# Temperature Scaling as a Free Lunch for *De Novo* Peptide Sequencing: Further Gains from Training-Free Candidate Rescoring

**DOI:** 10.64898/2026.11.23.768748

**Authors:** Peng Xiong, Yining Wang, Ruidi Zhang, Kai Wang, Hongtao Xu, Weiqi Wu, Weizun Zhao, Xugang Wang, Zheng Tang

**Affiliations:** Institute of AI for Industries, Chinese Academy of Sciences, China; School of Computer Science and Technology, University of Science and Technology of China, China; Jiangsu Key Laboratory of AI for Industries, Institute of AI for Industries, Chinese Academy of Sciences, China

## Abstract

*De novo* peptide sequencing infers peptide sequences directly from tandem mass spectra without relying on a reference database, enabling the identification of peptides absent from reference collections. Autoregressive models commonly decode with beam search and return the candidate with the highest model score. We first show that decoding temperature constitutes a free lunch for this pipeline: raising the decoding temperature alone improves top-1 peptide recall of Casanovo, InstaNovo, and ContraNovo by 0.5–1.0 percentage points without retraining, changes to model weights, or additional decoding passes. Temperature affects both the sequences admitted to the beam-search candidate pool and the final candidate scores, although the former effect depends on the implementation. Under the *T* = 1.0 baseline ranking defined here, the proportion of spectra whose candidate pools contain the correct peptide exceeds top-1 peptide recall by 9–12 percentage points, revealing additional ranking headroom. To exploit this headroom, we then introduce FragR (Fragment-based Rescoring), a training-free method that combines model scores with fragment evidence computed directly from raw spectral peaks: intensity-weighted fragment coverage and b/y-ion ladder evidence. On the nine-species V1 dataset (approximately 1.53 million monoisotopic spectra), combining an elevated decoding temperature with FragR improves peptide recall over the *T* = 1.0 baseline by 1.24–1.49 percentage points across the three models. The combined approach also improves recall on the nine-species V2 dataset (232,738 spectra), which includes isotope-shifted precursors. Asynchronous CPU rescoring overlaps with GPU decoding, so the additional FragR step adds little end-to-end inference overhead.

## 1 Introduction

Mass spectrometry is a central technology in bottom-up proteomics, supporting both protein identification and quantification [Aebersold and Mann, 2003]. Mainstream workflows rely on database searching: reference protein sequences are digested in silico according to enzyme-specific rules, theoretical fragment ions are generated for the resulting candidate peptides, and experimental spectra are matched and scored against these candidates [Eng et al., 1994, Perkins et al., 1999]. The best-scoring matches are typically filtered at a 1% false discovery rate (FDR) [Elias and Gygi, 2007], after which peptide assembly and protein inference yield proteome-level conclusions [Nesvizhskii et al., 2007]. This workflow presupposes a sufficiently complete and well-annotated reference proteome. Its applicability becomes limited when genomes are unsequenced or poorly annotated, or when target sequences—such as neoantigens and antibody variable regions—are absent from the reference database [Chong et al., 2020, Liao et al., 2024]. *De novo* peptide sequencing addresses this limitation by inferring the most likely amino acid sequence directly from an experimental spectrum, without a reference database. It is therefore valuable for discovering non-canonical and variant peptides, sequencing and assembling antibodies, and analyzing metaproteomes [Bandeira et al., 2008, Chong et al., 2020, Xiong et al., 2026, Van Den Bossche et al., 2025].

Experimental spectra frequently contain noise peaks, missing fragment signals, co-fragmented peptides, and side-chain neutral losses, making *de novo* sequencing a highly underdetermined inference problem. Classical methods, including Lutefisk [Taylor and Johnson, 2001], PEAKS [Ma et al., 2003], PepNovo [Frank and Pevzner, 2005], and Novor [Ma, 2015], combined handcrafted scoring functions with heuristic, probabilistic, or dynamic-programming-based sequence search. DeepNovo [Tran et al., 2017] introduced deep learning to this task and demonstrated substantial accuracy gains from data-driven modeling. PointNovo [Qiao et al., 2021] and PepNet [Liu et al., 2023] subsequently improved model architectures and the representation of high-resolution spectra. Casanovo [Yilmaz et al., 2022] reformulated the task as sequence-to-sequence translation, introduced a Transformer architecture, and used positional encodings with adjustable wavelengths to represent mass-to-charge values at the required precision. It has since become a common baseline paradigm, with subsequent methods broadly divided into autoregressive (AR) and non-autoregressive (NAR) decoding routes [Xia et al., 2025a].

Despite their architectural differences, these models ultimately return the candidate assigned the highest score under their respective decoding procedures. This decoding stage offers two complementary opportunities for improving accuracy without retraining. The first is temperature scaling, which divides decoder logits by a temperature before softmax normalization [Hinton et al., 2015]. Although temperature is often treated as a sampling-diversity parameter, it can affect both candidate generation and final ranking in beam-search decoders. Once the temperature has been selected, using it requires no model update, additional decoding pass, or post-processing component. We therefore call the resulting temperature-only improvement a *free lunch*.

The second opportunity arises after the candidate pool has been generated: the correct peptide may already be present but remain below rank one. Existing approaches to this selection problem generally require an additional ranker trained on labeled data or predictions from multiple models [Yang et al., 2019, Seo et al., 2025, Liao et al., 2024, Qiu et al., 2025]. To obtain further gains without training, we treat each beam-search candidate pool as a miniature, spectrum-specific database-search space. Selecting the sequence that best matches the spectrum is then analogous to peptide-spectrum match (PSM) scoring in data-dependent acquisition (DDA) database search [Eng et al., 1994, Craig and Beavis, 2004]. We introduce FragR as a complementary post-processing step that linearly combines the model score with b/y-ion ladder evidence, intensity-weighted fragment coverage, and precursor-mass consistency computed from raw peaks. Satellite (isotope) peaks are considered in ladder matching but are not matched when computing intensity-weighted coverage. Its CPU implementation overlaps asynchronously with GPU decoding, resulting in minimal additional latency. Across Casanovo, InstaNovo, and ContraNovo on approximately 1.53 million nine-species spectra, combining temperature scaling with FragR improves peptide recall over the *T* = 1.0 baseline by 1.24–1.49 percentage points. The combined approach also improves recall on spectra with isotope-shifted precursors. Section 2 reviews related work, Section 3 presents the empirical motivation, Section 4 describes the complementary rescoring method, Section 5 reports the experiments, Section 6 concludes the paper, and Section 7 outlines future work.

## 2 Related Work

### 2.1 Autoregressive *De Novo* Peptide Sequencing

Autoregressive methods generate amino-acid tokens sequentially, as exemplified by Casanovo [Yilmaz et al., 2022, 2024] and InstaNovo [Eloff et al., 2025]. At inference, beam search expands the explored sequence space under precursor-mass constraints: sequences whose cumulative mass exceeds the tolerance trigger early termination, and candidates outside the tolerance window are penalized during top-1 selection. Recent improvements fall into six broad directions. First, missing-fragment modeling and imputation include the two-stage graph model in GraphNovo [Mao et al., 2023], latent-space imputation in LIPNovo and LIPNovo+ [Du et al., 2025a], and error-guided fragment imputation in GyroNovo [Mekki et al., 2026]. Second, ContraNovo aligns spectral and sequence representations through contrastive learning [Jin et al., 2024]. Third, *π*-HelixNovo introduces complementary-spectrum encoding to incorporate additional ion information [Yang et al., 2024]. Fourth, bidirectional generation is explored by NovoB [Lee and Kim, 2024] and BiATNovo [Yang et al., 2023], while *π*-HelixNovo2 combines complementary-spectrum encoding with an ensemble of forward and reverse decoders [Yang et al., 2026]. Fifth, RNovA uses rotary positional embeddings to model spectral peaks and supports zero-shot prediction of post-translational modifications unseen during training [Mao et al., 2026]. Sixth, retrieval-augmented methods use training-derived reference information: SearchNovo retrieves similar PSMs and incorporates reference peptides through a fusion layer to supplement missing fragment evidence [Xia et al., 2025b], whereas ReNovo constructs a datastore from training data and queries it during autoregressive inference while retaining the ability to predict novel peptides [Chen et al., 2025a]. Both approaches require training-derived datastores and retrieval infrastructure; SearchNovo additionally trains a fusion module.

### 2.2 Non-Autoregressive *De Novo* Peptide Sequencing

Non-autoregressive methods predict amino-acid distributions at all positions in parallel, allowing each position to use bidirectional context. Decoding consequently shifts from prefix-based beam search to combinatorial optimization under precursor-mass constraints. *π*-PrimeNovo [Zhang et al., 2025a] removes the causal mask, trains with a connectionist temporal classification (CTC) loss [Graves et al., 2006], and uses knapsack-style dynamic programming to select the highest-scoring mass-compatible sequence. RefineNovo improves non-autoregressive training with curriculum learning to stabilize CTC training [Zhang et al., 2025b]. PowerNovo2 models the spectrum-to-peptide distribution with conditional normalizing flows, combining variational inference with tractable likelihood estimation and parallel decoding [Petrovskiy et al., 2026]. InstaNovo+ instead uses a diffusion model to iteratively refine an initial peptide prediction [Eloff et al., 2025].

### 2.3 Scaling Training Data and Species Coverage

Both decoding paradigms have also benefited from larger training corpora and broader species coverage. Casanovo achieved substantial gains after retraining on approximately 30 million PSMs from MassIVE-KB [Yilmaz et al., 2024, Wang et al., 2018]. MassNet constructed a spectral corpus from approximately 30 TB of raw data, covering 35 species and 558 million PSMs, and used non-autoregressive curriculum learning to train XuanjiNovo [A et al., 2026]. OmniNovo integrated MassIVE-KB with 20 PTM-enriched datasets to construct a corpus of 51.8 million PSMs [Chen et al., 2025b]. It represents 11 types of post-translational modification as separate vocabulary tokens and combines mass- and biochemistry-constrained decoding to sequence modified proteomes.

### 2.4 Temperature Scaling in Sequence Decoding

Temperature scaling divides decoder logits by *T* before applying softmax, thereby controlling the sharpness of the token distribution [Hinton et al., 2015]. In language-model generation, increasing the temperature can improve output diversity, whereas an excessively high temperature can reduce sample quality. Recent work on multi-sample inference further shows that the best temperature depends on the model, task, and aggregation strategy, and proposes entropy-based criteria for automatic temperature selection [Du et al., 2025b]. These findings motivate temperature as an inference-time control, but results for stochastic sampling and multi-sample aggregation do not directly characterize beam-search decoding in *de novo* sequencing.

### 2.5 Candidate Reranking and Rescoring

An orthogonal line of work improves candidate selection without changing the underlying sequence generator. This idea can be traced to pNovo 3 [Yang et al., 2019], which generates candidates by spectrum-based dynamic programming, derives features from similarities between pDeep-predicted and experimental spectra [Zhou et al., 2017], and trains an SVM-rank model for reranking. Its pipeline is coupled to the pNovo candidate generator, limiting its use as generic post-processing. NovoRank [Seo et al., 2025] combines spectral clustering based on spectral similarity, precursor *m/z*, and retention time with a learned ranker to refine candidates from multiple tools; it requires labeled training data and retention-time information. MARS [Liao et al., 2024] targets immunopeptidomics and combines average local confidence, MHC-binding rank, and retention time for candidate selection. RankNovo [Qiu et al., 2025] stacks candidates from six base models—three architectures and their reverse-decoding variants—as multiple sequence alignments and uses axial attention for deep cross-model reranking. It reports state-of-the-art peptide recall on the nine-species V1 benchmark, but its inference cost grows approximately linearly with the number of base models. These methods therefore require trained components, multiple model outputs, or application-specific dependencies (Table 1).

**Table 1:**
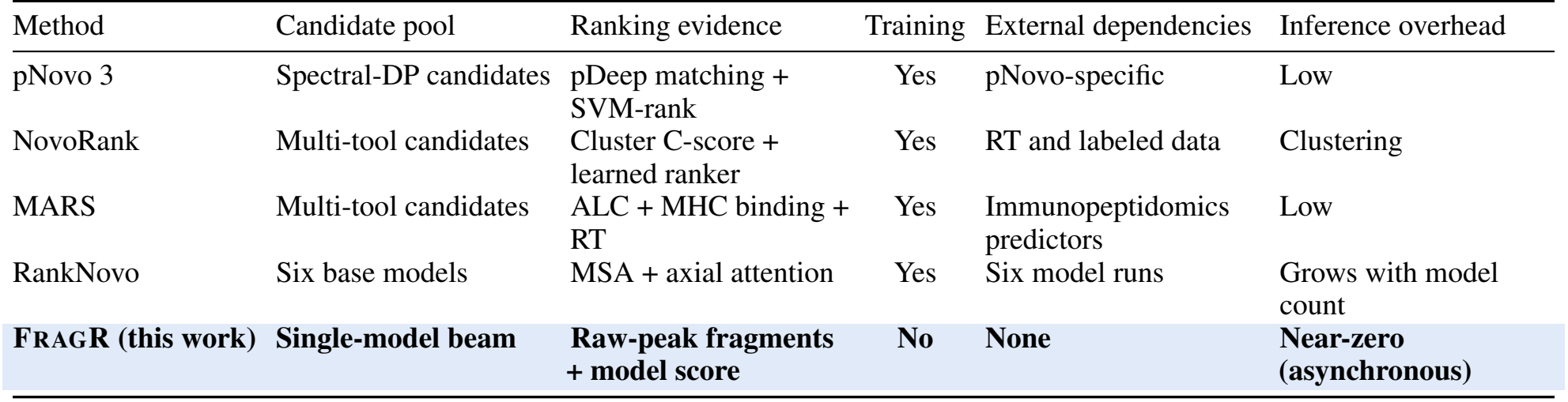
Comparison of candidate-reranking methods. The FragR row is highlighted in light blue.

| Method | Candidate pool | Ranking evidence | Training | External dependencies | Inference overhead |
|---|---|---|---|---|---|
| pNovo 3 | Spectral-DP candidates | pDeep matching + SVM-rank | Yes | pNovo-specific | Low |
| NovoRank | Multi-tool candidates | Cluster C-score + learned ranker | Yes | RT and labeled data | Clustering |
| MARS | Multi-tool candidates | ALC + MHC binding + RT | Yes | Immunopeptidomics predictors | Low |
| RankNovo | Six base models | MSA + axial attention | Yes | Six model runs | Grows with model count |
| **FRAGR (this work)** | **Single-model beam** | **Raw-peak fragments + model score** | **No** | **None** | **Near-zero (asynchronous)** |

A concurrent benchmark evaluated eight models and three post-processing tools: Spectralis [Klaproth-Andrade et al., 2024], InstaNovo+ [Eloff et al., 2025], and MS^2^Rescore [Declercq et al., 2022]. It distinguished confidence rescoring, evaluated by the area under the precision–coverage (PC) curve, from candidate selection, in which reranking changes the top-1 prediction [van Puyenbroeck et al., 2026]. The rescoring experiments showed no consistent improvement over native model scores. Candidate-selection results were mixed: Spectralis and InstaNovo+ did not improve Casanovo’s ranking, whereas MS^2^Rescore variants improved selection in some settings, particularly when the gold-standard candidate had been appended to an otherwise incomplete candidate list. The study also associated many errors with insufficient fragment coverage across models. Importantly, PC-area measurements quantify confidence calibration and filtering rather than fixed-pool top-1 selection, and its candidate-selection experiments focused on Casanovo without varying decoding temperature. Our study complements this benchmark by evaluating temperature scaling as a direct decoding intervention and reranking against same-temperature baselines, providing direct evidence for both settings. FragR occupies a distinct operating point: a single-model beam-search pool, training-free classical fragment evidence, and near-zero additional inference latency.

## 3 Empirical Motivation: Temperature Scaling and Ranking Headroom

### 3.1 Mismatches Between Decoding Settings and Benchmark Data

The nine-species dataset (V1), introduced by DeepNovo [Tran et al., 2017], contains approximately 1.53 million spectra and is one of the most widely used benchmarks in subsequent work. Some reported results for autoregressive models may be affected by a mismatch between inference configurations and dataset characteristics. First, the default decoding configuration of Casanovo and related autoregressive models sets the minimum peptide length to six, whereas V1 contains 7,178 spectra (0.47%) whose ground-truth peptides have length five. For these spectra, the correct peptide is pruned during decoding and cannot be recovered by any subsequent selection procedure. Second, the default configuration permits a +1 isotope error, allowing a candidate mass to differ from the precursor by approximately 1.003 Da, whereas all V1 precursors are monoisotopic and contain no isotope-shift annotation. A top-1 candidate selected under the shifted-isotope hypothesis therefore differs from the ground truth by approximately 1 Da and is necessarily incorrect on V1. In contrast, non-autoregressive decoders such as *π*-PrimeNovo impose no minimum-length constraint and use the hard mass constraint in Eq. (1), without an isotope hypothesis [Zhang et al., 2025a]. This configuration is better matched to monoisotopic data. Consequently, default settings can underestimate the performance of autoregressive models on this benchmark.

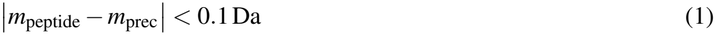

### 3.2 Candidate Pools, Oracle Recall, and Error Categories

Beam search produces a candidate pool for each spectrum, consisting of completed sequences that either emit an end token or exceed the precursor-mass tolerance during generation. The model ordinarily returns the candidate with the highest score. We partition all spectra into four mutually exclusive and exhaustive categories according to the top-1 prediction and the contents of the candidate pool: C1, the top-1 prediction has an isotope-assignment error (an *isotope trap*) and the pool contains the correct answer; C2, the top-1 prediction is incorrect and the pool contains the correct answer; C3, the top-1 prediction is correct; and C4, the pool does not contain the correct answer (a *search failure*). No post-processing method that leaves generation unchanged can recover spectra in C4. The sum C1+C2+C3 is the proportion of spectra whose pools contain the correct answer, which we call the oracle, and C4 = 1− oracle. Correctness follows the exact peptide-match criterion defined in Section 5.1.

As shown in Fig. 1a, with beam width 10, minimum peptide length five (no minimum-length setting is used for InstaNovo), and a maximum isotope error of one, the three baseline models achieve oracle values of 67.4–72.7%. These values exceed the nine-species V1 state-of-the-art top-1 recall of 66.0% reported for RankNovo [Qiu et al., 2025]. The oracle is an upper bound rather than a directly comparable top-1 result; it quantifies the headroom available to candidate reranking. Our rescoring objective is to recover correct answers from C1+C2 by promoting them to rank one, after temperature scaling has been applied within the decoder. We also define recall@*k* as the proportion of spectra for which the correct answer occurs within the top *k* candidates. This quantity increases monotonically with *k* and reaches the oracle at *k* = 10 (Fig. 1b).

**Figure 1:**
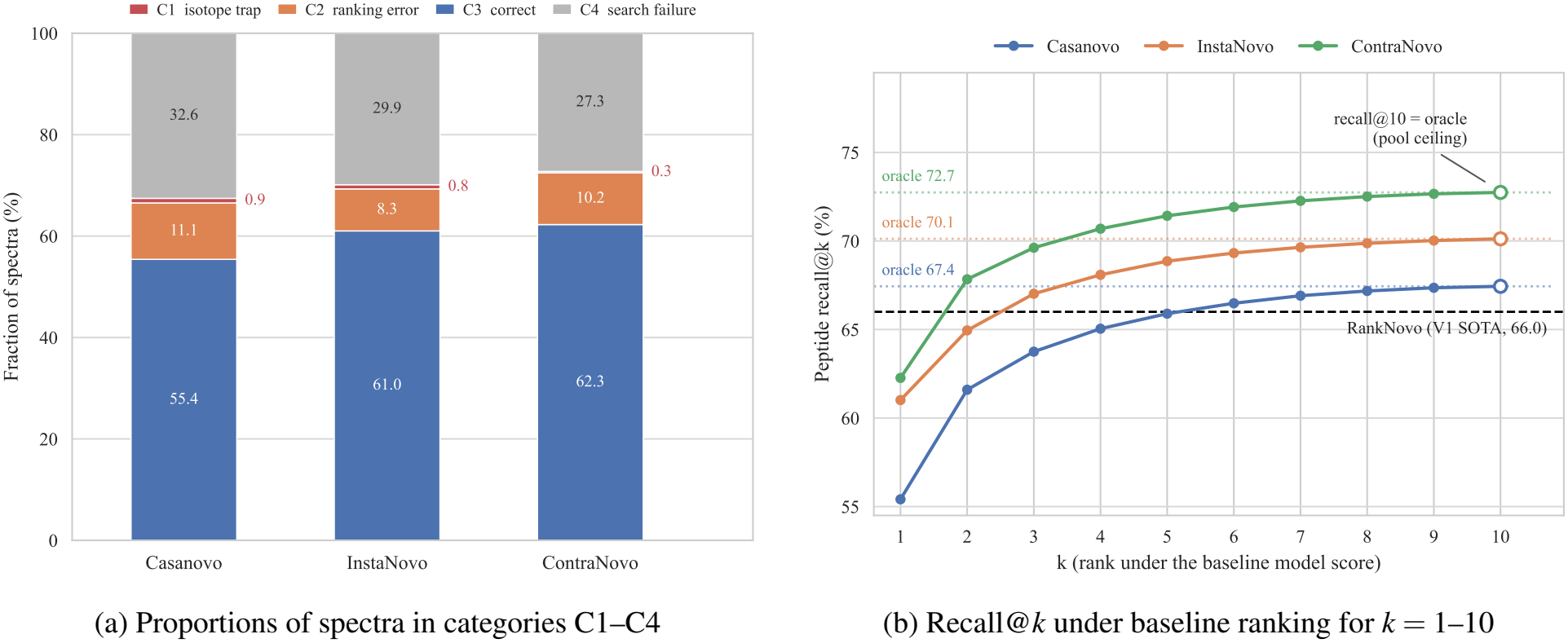
Error decomposition and recall curves for baseline beam-search candidate pools. The black dashed line marks the V1 state-of-the-art recall of 66.0% reported by RankNovo.

### 3.3 Temperature Scaling as a Free Lunch

Temperature scaling divides each decoder logit by *T* before softmax normalization (Appendix B.4). Increasing *T* flattens the token distribution, whereas decreasing *T* sharpens it [Hinton et al., 2015]. Its effects on generation diversity and downstream accuracy depend on the decoding procedure and task [Du et al., 2025b]. We use *free lunch* specifically for the temperature-only intervention: once *T* has been selected, inference uses the same trained model and beam-search pipeline, without an additional decoding pass or post-processing component.

Temperature affects beam search through two routes in the implementations studied here. First, it can change the sequences admitted to the candidate pool: the pools produced by InstaNovo and ContraNovo vary with temperature, whereas those produced by Casanovo do not (Appendix B). Second, it changes the final candidate scores and can therefore alter their ranking. The first route changes the oracle, whereas the second can improve top-1 recall even when the candidate pool remains fixed.

We scanned temperatures from 0.2 to 2.0 on the nine-species V1 benchmark. Temperatures above one consistently improved recall without adding a decoding pass. For ContraNovo, increasing the temperature from 1.0 to 2.0 improved the model’s own top-1 recall by approximately 1.0 percentage points and increased the candidate-pool oracle from 72.7% to 73.6% (Fig. 2). Temperature scaling therefore provides a direct gain for all three models and, for InstaNovo and ContraNovo, also expands the C1+C2 headroom available to subsequent candidate reranking.

**Figure 2:**
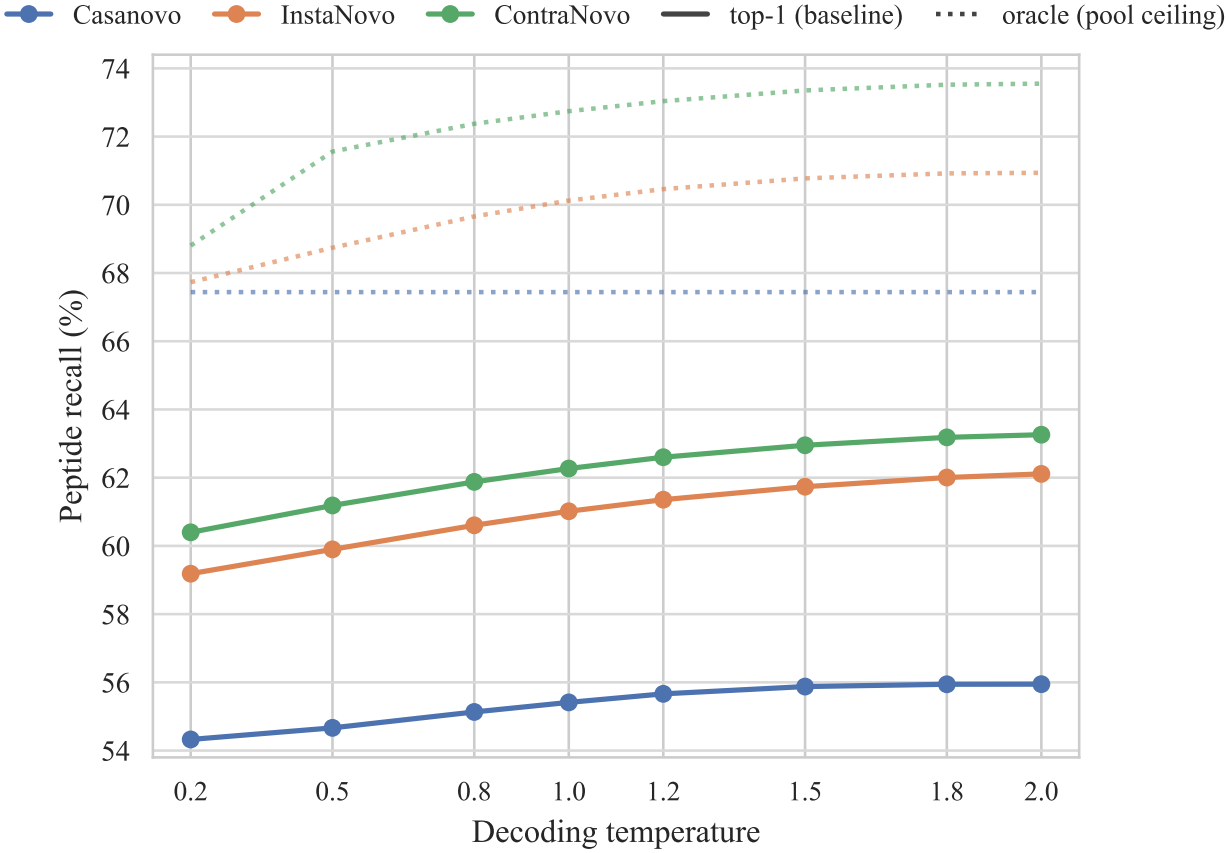
Effect of decoding temperature on baseline top-1 peptide recall and the candidate-pool upper bound. Solid lines show top-1 recall under baseline ranking, and dotted lines show the corresponding oracle values.

## 4 FragR: Fragment-Based Candidate Rescoring

Temperature scaling operates within each model’s native decoder. FragR is a complementary post-processing step applied to the candidate pool produced at any chosen temperature; it changes neither the model weights nor the pool itself. For each spectrum *s*, represented by its peak list, precursor *m/z*, and charge, beam search produces a candidate pool *ℬ*(*s*). The baseline ranking returns the candidate with the highest model score. FragR instead treats *ℬ*(*s*) as a spectrum-specific miniature database-search space and assigns each candidate *c* ∈ *ℬ*(*s*) a score that combines model confidence with fragment evidence:

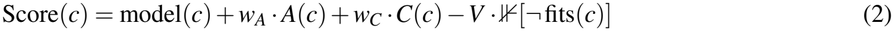

Here, model(*c*) is the model-specific candidate score used in both baseline ranking and rescoring; implementation differences and score alignment are described in Appendix B. The terms *A* and *C* are two fragment-evidence scores computed from unprocessed raw spectral peaks, as defined below. The predicate fits(*c*) tests precursor-mass consistency: a candidate is accepted if its neutral mass agrees with the precursor within *±*50 ppm under either the monoisotopic or the configured maximum isotope tolerance. Casanovo and ContraNovo natively penalize mass-inconsistent candidates with a constant −1 term, whereas InstaNovo subtracts 10^6^. In the aligned model scores (Appendix B.3), we remove the native penalties and adopt the *V* = 10^6^ convention of InstaNovo [Eloff et al., 2025], which guarantees that every mass-consistent candidate ranks above all candidates that violate the constraint.

The two evidence terms are motivated by fragment-match counts and matched intensities used in classical database-search engines such as SEQUEST [Eng et al., 1994] and X!Tandem [Craig and Beavis, 2004]. FragR transfers this evidence to beam-search candidate pools without training. The coefficients are fixed to *w*_*A*_ = *w*_*C*_ = 0.25 across all models, temperatures, and datasets; the procedure contains no trainable parameters. Both evidence terms lie in [0, 1], giving a maximum additive reward of *w*_*A*_ + *w*_*C*_ = 0.5. This scale is compatible with observed within-pool model-score differences: on V1 at *T* = 1.0, 90% of spectra have top-1–to–top-2 differences no greater than 0.3.

For a candidate *c* of length *L*, we generate the theoretical *m/z* values of *b*_1_, …, *b*_*L*−1_ and *y*_1_, …, *y*_*L*−1_ at each charge state *z* ∈ {1, …, *z*_max_} using standard b/y-ion formulas. Here, *z*_max_ = max(1, *z*_*p*_−1), where *z*_*p*_ is the precursor charge, giving *n*_ions_ = 2(*L*− 1)*z*_max_ theoretical ions. A theoretical ion is matched if the raw spectrum contains a peak within 20 ppm of its *m/z* value. For ladder evidence, the matching procedure also considers satellite (isotope) peaks associated with these theoretical ions. A cleavage site is counted as matched if any of its charge states is matched; there are *n*_pos_ = 2(*L*− 1) cleavage positions across the b- and y-ion series. Let *n*_*b*_ and *n*_*y*_ denote the numbers of matched cleavage sites on the b and y sides, respectively. The ladder evidence is

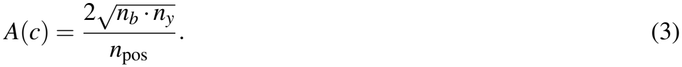

The geometric mean requires supporting evidence from both ion series.

The second term incorporates peak intensity to help distinguish signal matches from noise matches. Let the relative intensity of raw peak *j* be *r* _*j*_ = *I*_*j*_*/* max_*k*_ *I*_*k*_. Intensity-weighted fragment coverage is defined as

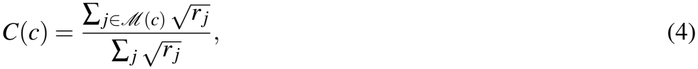

where *ℳ* (*c*) is the set of raw peaks matched directly to any theoretical b/y ion of candidate *c*. Satellite (isotope) peaks are not matched for this intensity term. Each peak is counted once regardless of how many theoretical ions it matches, and the denominator sums over all raw peaks in the spectrum. Thus, *C* ∈ [0, 1] measures the candidate’s share of the total square-root-transformed relative intensity, 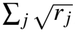, rather than its share of the raw intensity. The square-root transformation compresses the dynamic range so that the base peak does not dominate the score.

Rescoring consists entirely of CPU numerical operations and has no cross-spectrum dependencies. Executing it synchronously would serialize candidate export and scoring within the inference loop. FragR therefore uses an asynchronous pipeline: a GPU producer process performs beam search in batches and publishes completed candidates to a shared-memory buffer, while a CPU consumer process rescores, sorts, and writes the candidates without blocking the producer. In steady state, production and consumption overlap, and end-to-end wall-clock time is close to that of decoding alone. We implemented the same interface in the inference repositories of all three baseline models, allowing the rescored output to replace the original top-1 prediction.

## 5 Experimental Evaluation

### 5.1 Datasets and Evaluation Metrics

Experiments were conducted on two datasets derived from the nine-species benchmark. V1 is the full monoisotopic dataset [Tran et al., 2017], containing 1,528,127 spectra; every precursor is reported as monoisotopic, with no isotope-shift annotation. V2 is a file-level subset of the revised nine-species benchmark [Wen and Noble, 2024]—commonly referred to as the nine-species V2 data—selected to include isotope-shifted spectra. It contains 232,738 spectra, of which 17,763 (7.6%) have precursors reported as M+1 isotope masses. We denote the ordinary-precursor and isotope-precursor subsets as e0 and e1, respectively. Files were selected to balance isotope-spectrum counts across species: 77 complete MGF files were chosen (listed in Table A3), yielding approximately 1,600 isotope spectra for eight species (range: 1,581–1,755). Yeast is the only marked deviation from this balance because its single available file contains 4,358 isotope spectra. All reported metrics are macro-averaged with equal weight across the nine species; V1 uses the full dataset.

Evaluation follows standard *de novo* sequencing conventions, with matching rules consistent with DeepNovo [Tran et al., 2017] and Casanovo [Yilmaz et al., 2022]. Given a predicted peptide 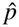 and a ground-truth peptide *p*, a two-pointer procedure aligns residues from the N terminus. A position can be paired only if the cumulative mass difference between the aligned portions is less than 0.5 Da, and a paired position is counted as matched only if the residue-mass difference is less than 0.1 Da. If the prefix alignment does not cover all positions, suffix alignment proceeds in the reverse direction from the C terminus, and the two alignment results are merged. Residues with identical masses or mass differences within the tolerance are treated as equivalent. The primary metric is peptide recall:

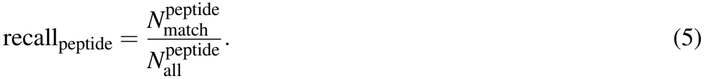

Here, 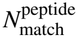 is the number of exactly correct peptides. A correct prediction must have the same length as the ground truth and, after the bidirectional alignment described above, every position must have a residue-mass difference below 0.1 Da while the cumulative mass difference remains below 0.5 Da. The denominator 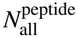 is the total number of evaluated spectra, including spectra with empty predictions, which are always counted as unrecalled.

### 5.2 Temperature Scaling and Further Gains from Rescoring

We evaluate eight temperatures from 0.2 to 2.0 for each model. Throughout the experiments, Casanovo refers to the model with weights trained on MassIVE data [Yilmaz et al., 2024], corresponding to code release v4.0.0 (see Code Availability). The baseline curves across temperatures measure the temperature-only effect. At each temperature, FragR is compared with the baseline ranking defined here on the same candidate pool, which isolates its additional effect. For ContraNovo, this baseline uses mean ordinary softmax probabilities; the original implementation’s second softmax during final ranking is removed for both baseline ranking and rescoring (Appendix B.3). Table 2 summarizes the *T* = 1.0 baseline and a common elevated setting of *T* = 1.8; Fig. 3 reports the complete temperature response.

**Table 2:**
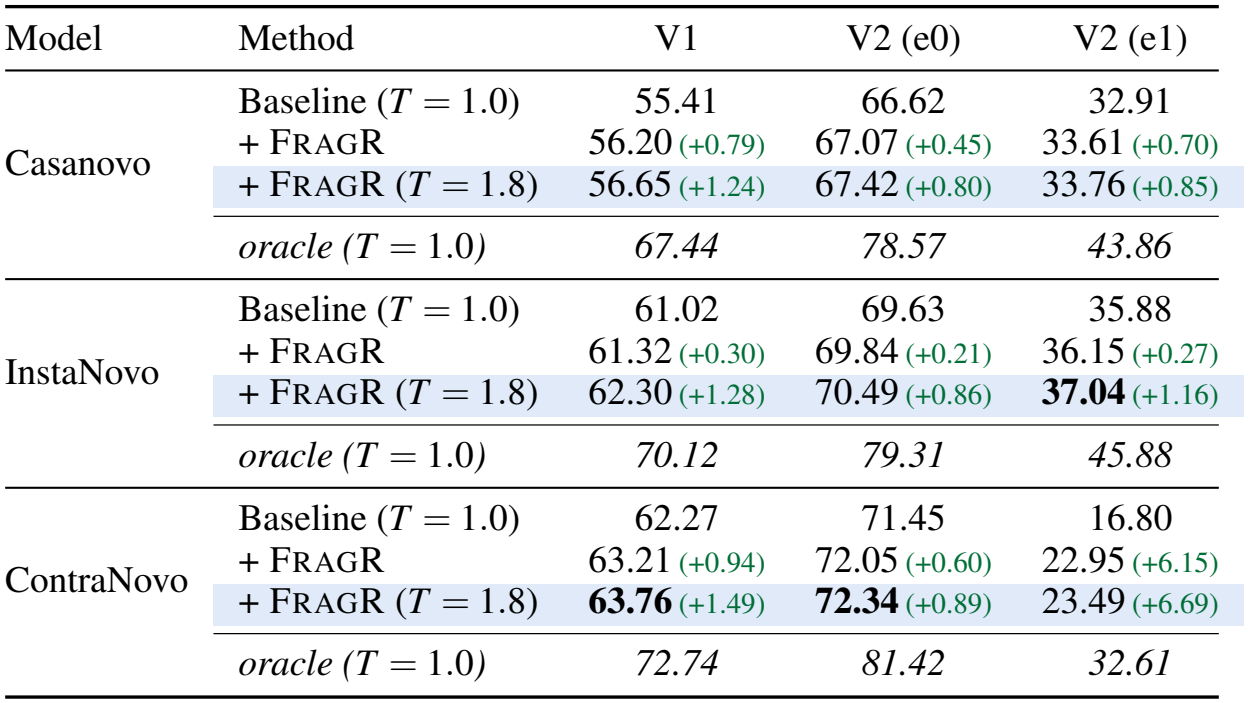
Combined effects of temperature scaling and FragR on nine-species macro-averaged peptide recall (%). “Baseline” denotes the *T* = 1.0 baseline ranking defined in Appendix B.3; “+ FragR” denotes rescoring at *T* = 1.0; and “+ FragR (*T* = 1.8)” combines temperature scaling with rescoring and is highlighted in light blue. Parentheses contain gains relative to the *T* = 1.0 baseline for the same dataset, in percentage points; the final setting therefore measures the combined gain, not the isolated effect of rescoring at *T* = 1.8. Positive and negative gains are green and red, respectively. The best value in each column is bold, and oracle denotes the candidate-pool upper bound at *T* = 1.0.

| Model | Method | V1 | V2 (e0) | V2 (e1) |
|---|---|---|---|---|
| Casanovo | Baseline (*T* = 1.0) | 55.41 | 66.62 | 32.91 |
|  | + FragR | 56.20 (+0.79) | 67.07 (+0.45) | 33.61 (+0.70) |
|  | + FragR (*T* = 1.8) | 56.65 (+1.24) | 67.42 (+0.80) | 33.76 (+0.85) |
|  | *oracle (T = 1.0)* | *67.44* | *78.57* | *43.86* |
| InstaNovo | Baseline (*T* = 1.0) | 61.02 | 69.63 | 35.88 |
|  | + FragR | 61.32 (+0.30) | 69.84 (+0.21) | 36.15 (+0.27) |
|  | + FragR (*T* = 1.8) | 62.30 (+1.28) | 70.49 (+0.86) | **37.04** (+1.16) |
|  | *oracle (T = 1.0)* | *70.12* | *79.31* | *45.88* |
| ContraNovo | Baseline (*T* = 1.0) | 62.27 | 71.45 | 16.80 |
|  | + FragR | 63.21 (+0.94) | 72.05 (+0.60) | 22.95 (+6.15) |
|  | + FragR (*T* = 1.8) | **63.76** (+1.49) | **72.34** (+0.89) | 23.49 (+6.69) |
|  | *oracle (T = 1.0)* | *72.74* | *81.42* | *32.61* |

**Figure 3:**
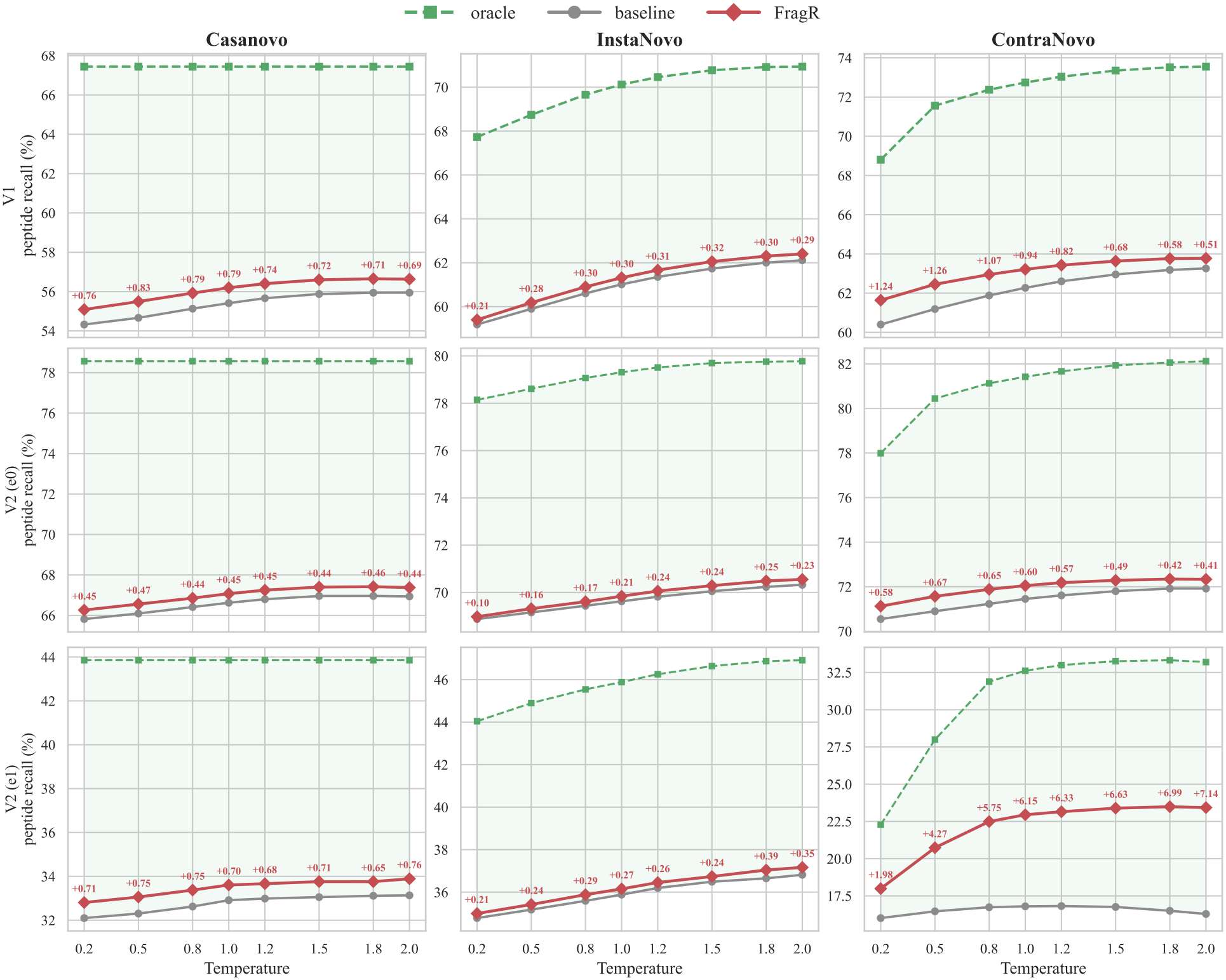
Peptide recall as a function of decoding temperature before and after rescoring. Each panel contains three curves: baseline (gray circles), FragR (red diamonds), and oracle (green dashed squares). Red numbers indicate the gain from FragR over the baseline at the same temperature, in percentage points. The oracle values in Table 2 are at *T* = 1.0; the oracle curves here instead use the candidate pool at each plotted temperature.

At *T* = 1.0, FragR improves V1 peptide recall for Casanovo, InstaNovo, and ContraNovo by 0.79, 0.30, and 0.94 percentage points, respectively. Both V2 subsets show gains in the same direction: +0.45/+0.21/+0.60 on e0 and +0.70/+0.27/+6.15 on e1, in the same model order. Combining *T* = 1.8 with FragR yields combined V1 gains over the *T* = 1.0 baseline of 1.24, 1.28, and 1.49 percentage points, respectively. The corresponding combined gains range from 0.80 to 0.89 percentage points on e0 and from 0.85 to 6.69 percentage points on e1; the isotope subset is analyzed further in Section 5.3. Across all model–species pairs on V1, these combined gains range from +0.46 to +2.39 percentage points, and all nine species show positive gains (Table A4).

The two interventions play distinct but complementary roles. Temperature scaling improves baseline top-1 recall and, for InstaNovo and ContraNovo, can also expand the candidate pool (Section 3.3). FragR then improves ranking at a fixed temperature, whether the correct candidate was already present or was newly admitted by temperature scaling.

Figure 3 compares all eight temperatures across three models and three data slices. The gray baseline curves show the recall at each temperature under baseline ranking, whereas the difference between the red and gray curves at the same temperature isolates the additional gain from FragR. At every model–dataset–temperature combination, the macro-averaged recall of FragR is no lower than that of the same-temperature baseline. Casanovo’s oracle curve is flat because its candidate pool does not change with temperature; its additional FragR gain is nearly temperature-invariant, ranging from 0.69 to 0.83 percentage points on V1. For ContraNovo, the additional V1 gain narrows from 1.24 percentage points at *T* = 0.2 to 0.51 percentage points at *T* = 2.0 because temperature improves the baseline more rapidly. On e1, by contrast, the additional gain expands from 1.98 to 7.14 percentage points, while the oracle increases from 22.3% to approximately 33.2%; rescoring converts much of this temperature-induced headroom into top-1 recall. InstaNovo shows moderate additional gains that increase slightly with temperature.

On V1, FragR gains over the same-temperature baseline are positive for every species at every temperature. On e0, the only negative species-level value is InstaNovo on honeybee at *T* = 1.5 (−0.03 percentage points, approximately two spectra). All other negative values occur on e1: InstaNovo on clambacteria is negative at five temperatures (*T* = 0.8, 1.0, 1.2, 1.5, 1.8), with a minimum of −0.23 percentage points; InstaNovo on bacillus (*T* = 0.2, −0.13 percentage points) and honeybee (*T* = 1.5, −0.06 percentage points) each has one negative value; and Casanovo on bacillus has one at *T* = 1.8 (−0.19 percentage points). Most e1 species contain approximately 1,600 spectra, except yeast with 4,358, so −0.23 percentage points corresponds to only about four spectra for a typical species. The numbers of spectra corrected and corrupted are analyzed in Section 5.4 and Fig. 4.

**Figure 4:**
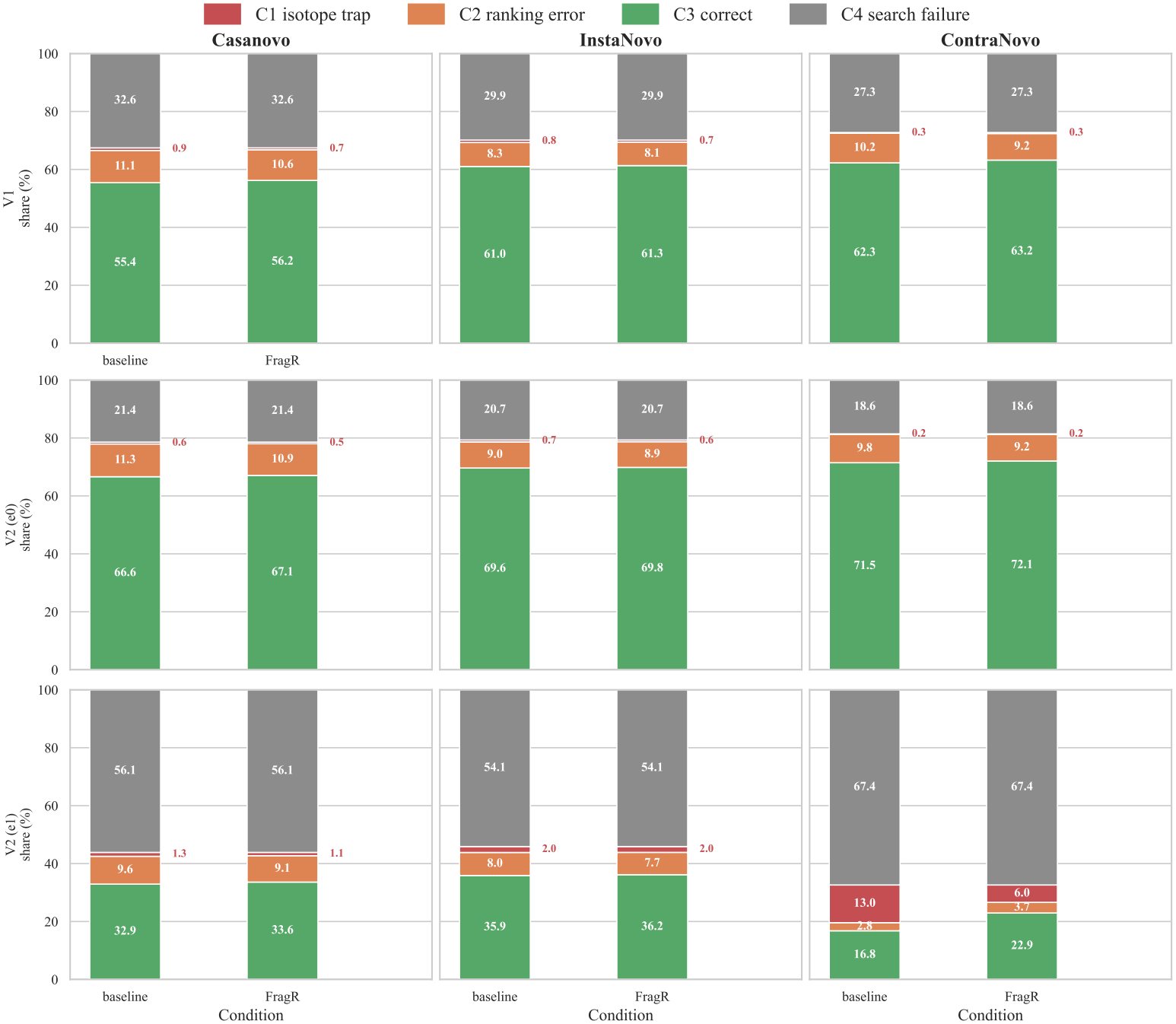
C1–C4 error decomposition before and after rescoring. Each panel contains two 100% stacked bars: baseline on the left and rescoring on the right. From bottom to top, the four segments are C3 (correct), C2 (ranking error), C1 (isotope trap), and C4 (search failure). C1 values are annotated beside the bars.

### 5.3 Results on Ordinary and Isotope-Shifted Precursors

The e0 subset serves as a consistency check. At *T* = 1.0, the three baseline recalls are 66.62%, 69.63%, and 71.45%, and FragR improves them by 0.45, 0.21, and 0.60 percentage points, respectively—gains of the same magnitude as on V1. Combining FragR with a common temperature of *T* = 1.8 yields combined gains over the *T* = 1.0 baseline of +0.80, +0.86, and +0.89 percentage points, respectively. The e0 oracle ranges from 78.6% to 81.4%. Evaluating this subset separately therefore does not change the qualitative complementarity between temperature scaling and rescoring.

The e1 subset (17,763 spectra, 7.6%) is the principal focus of the V2 evaluation. All baseline models degrade sharply on this subset: Casanovo and InstaNovo fall from 66.6% and 69.6% on e0 to 32.9% and 35.9% on e1, respectively, while ContraNovo drops from 71.5% to 16.8%. On e1, the reported precursor mass is approximately 1 Da above the actual peptide mass (Section 3.1), which interferes with mass-conditioned decoding. The additional FragR gain is especially large for ContraNovo: at *T* = 1.0, the three models improve by 0.70, 0.27, and 6.15 percentage points, respectively. ContraNovo recovers to 22.95% with FragR at *T* = 1.0 and reaches 23.49% when rescoring is combined with *T* = 1.8. At the species level (Table A5), ContraNovo improves on all nine e1 species, with FragR gains at *T* = 1.0 ranging from +3.30 percentage points on clambacteria to +9.55 percentage points on yeast; the largest combined gain at *T* = 1.8 reaches +11.11 percentage points. At *T* = 1.0, species-level gains on e1 range from +0.29 to +1.28 percentage points for Casanovo and from −0.06 to +0.58 percentage points for InstaNovo. At the macro level, the FragR gain over the same-temperature baseline is positive across all 24 model–temperature combinations on e0 and across all 24 on e1.

### 5.4 Ranking Changes and Error Transitions

Figure A1 compares recall@*k* before and after rescoring at several cutoff depths *k*. At *k* = 2, the three models improve over baseline ranking by 0.53, 0.16, and 0.73 percentage points, respectively; at *k* = 5, the corresponding gains are 0.14, 0.05, and 0.46 percentage points. The gains decrease monotonically as *k* increases, and at *k* = 10 both rankings coincide with the oracle. Thus, rescoring changes the depth distribution of correct answers without changing the candidate pool. Under baseline ranking, recall@2 exceeds recall@1 by 6.2, 3.9, and 5.6 percentage points for the three models, respectively.

Table 3 reports the numbers of spectra corrected and corrupted for each species at *T* = 1.0. On V1, the three models correct 17,884, 6,623, and 20,901 spectra and corrupt 5,898, 2,274, and 6,383 spectra, respectively. The correction-to-corruption ratio remains between 2.9 and 3.3:1, the corruption rate does not exceed 0.42%, and corrections outnumber corruptions for all nine species. The ratios are similar on the ordinary-precursor V2 subset e0 (2.9–3.9:1). The isotope subset e1 exhibits a pronounced asymmetry: ContraNovo corrects 1,242 spectra and corrupts only 57 (21.8:1). This asymmetry also holds at the species level; for example, the corresponding counts are 433 versus 17 on yeast and 60 versus 2 on clambacteria. InstaNovo shows the same pattern at a smaller scale (87 corrections versus 32 corruptions). On clambacteria, however, it corrects five spectra and corrupts six, a net loss of one spectrum, consistent with the −0.06 percentage points change at *T* = 1.0 in Table A5.

**Table 3:** Per-spectrum changes caused by rescoring at *T* = 1.0 (numbers of spectra). “Corrected” means that the baseline top-1 prediction is incorrect but becomes correct after rescoring; “corrupted” means the reverse. The last column gives the correction-to-corruption ratio; clambact. abbreviates clambacteria.

| Model | Dataset |  | bacillus | clambact. | honeybee | human | mmazei | mouse | ricebean | tomato | yeast | Total | Ratio |
|---|---|---|---|---|---|---|---|---|---|---|---|---|---|
| Casanovo | V1 | Corrected | 3,755 | 1,539 | 3,678 | 1,503 | 2,023 | 272 | 595 | 3,080 | 1,439 | 17,884 | 3.0 |
|  |  | Corrupted | 1,317 | 598 | 1,267 | 459 | 742 | 134 | 167 | 881 | 333 | 5,898 |  |
|  | V2 (e0) | Corrected | 110 | 134 | 54 | 98 | 141 | 68 | 1,261 | 114 | 116 | 2,096 | 2.9 |
|  |  | Corrupted | 58 | 43 | 35 | 45 | 53 | 56 | 364 | 42 | 33 | 729 |  |
|  | V2 (e1) | Corrected | 14 | 19 | 25 | 19 | 21 | 18 | 17 | 8 | 82 | 223 | 2.7 |
|  |  | Corrupted | 9 | 7 | 9 | 14 | 6 | 6 | 4 | 3 | 26 | 84 |  |
| InstaNovo | V1 | Corrected | 1,485 | 574 | 1,291 | 602 | 764 | 107 | 262 | 1,024 | 514 | 6,623 | 2.9 |
|  |  | Corrupted | 511 | 178 | 495 | 212 | 283 | 40 | 58 | 339 | 158 | 2,274 |  |
|  | V2 (e0) | Corrected | 48 | 49 | 23 | 48 | 63 | 44 | 431 | 60 | 44 | 810 | 3.0 |
|  |  | Corrupted | 14 | 18 | 17 | 10 | 25 | 15 | 140 | 13 | 16 | 268 |  |
|  | V2 (e1) | Corrected | 3 | 5 | 7 | 14 | 10 | 6 | 5 | 3 | 34 | 87 | 2.7 |
|  |  | Corrupted | 3 | 6 | 1 | 4 | 5 | 0 | 1 | 1 | 11 | 32 |  |
| ContraNovo | V1 | Corrected | 4,547 | 1,791 | 4,879 | 1,672 | 2,179 | 324 | 711 | 3,132 | 1,666 | 20,901 | 3.3 |
|  |  | Corrupted | 1,362 | 556 | 1,452 | 689 | 677 | 129 | 134 | 810 | 574 | 6,383 |  |
|  | V2 (e0) | Corrected | 162 | 161 | 79 | 111 | 145 | 94 | 1,190 | 86 | 122 | 2,150 | 3.9 |
|  |  | Corrupted | 60 | 42 | 34 | 61 | 53 | 41 | 217 | 20 | 23 | 551 |  |
|  | V2 (e1) | Corrected | 71 | 60 | 131 | 169 | 80 | 129 | 60 | 109 | 433 | 1,242 | 21.8 |
|  |  | Corrupted | 7 | 2 | 5 | 12 | 4 | 4 | 3 | 3 | 17 | 57 |  |

Figure 4 decomposes the errors at *T* = 1.0 into the four categories defined in Section 3.2. On V1, the residual headroom after rescoring, defined as the difference between the oracle and top-1 recall, is 8.8–11.2 percentage points. C2 (ranking errors) accounts for 8.1–10.6 percentage points, or more than 90% of this headroom, whereas C1 (isotope traps) accounts for only 0.35–0.68% of all spectra, macro-averaged across the nine species. C4 (search failures) accounts for 27–33% of all spectra and reflects a limitation of generation rather than candidate selection. The isotope subset has a different composition. For e1 spectra, the precursor is reported as M+1, so the correct candidate should be scored under the +1 isotope hypothesis. Here, C1 means that the baseline ranks a monoisotopic candidate first—the opposite mismatch direction from V1, where the data are monoisotopic but the top-1 prediction incorrectly uses an isotope-shifted candidate. For ContraNovo on e1, C1 decreases from 13.0% to 6.0%, while C3 increases from 16.8% to 22.9%. Thus, the corrections in Table 3 arise primarily from resolving this isotope-role mismatch. Together with temperature scaling, FragR recovers 10–14% of the baseline headroom (9.1–12.0 percentage points); the remainder is still present in the candidate pool.

### 5.5 Ablation Study and Weight Sensitivity

Figure A2 ablates the two evidence terms in Eq. (2). On V1 at *T* = 1.0, the ladder term *A* alone provides 80–91% of the full gain: the three models improve by +0.65, +0.24, and +0.86 percentage points, respectively, compared with +0.79, +0.30, and +0.94 percentage points for the complete score. The marginal contribution of the intensity-coverage term *C* is +0.06 to +0.14 percentage points. Its contribution is more pronounced in the isotope setting: for ContraNovo on e1, the gain increases from +5.47 percentage points with *A* alone to +6.15 percentage points with the complete score, a marginal contribution of +0.68 percentage points. Across the full temperature range, combining both terms yields positive gains.

Figure A3 shows a grid scan over the weights *w*_*A*_, *w*_*C*_ ∈{0, 0.25, 0.5, 1}; the horizontal axis in the figure is labeled 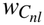. The frozen setting (0.25, 0.25) lies on a plateau of positive gains and is only 0.04–0.23 percentage points below the best setting in the grid. At *w*_*A*_ = 1, performance drops sharply: recall falls below the baseline for Casanovo and ContraNovo, by up to 0.24 and 0.86 percentage points, respectively. InstaNovo does not exhibit this decline.

### 5.6 Inference Overhead and Asynchronous Execution

We deployed FragR as an asynchronous pipeline in the inference repositories of all three models. Inference throughput was measured on the nine-species V1 data for both the baseline and the asynchronous pipeline, and end-to-end runtime was decomposed into GPU decoding, candidate publication, and CPU rescoring. Exporting candidate pools synchronously inside the inference loop reduces throughput by 3–7%. The asynchronous pipeline moves candidate export and rescoring to CPU processes that run concurrently with GPU decoding. End-to-end wall-clock time is consequently close to GPU decoding time, exceeding it by 0.5–4.5%, with the difference diminishing as batch size increases. Rescoring therefore adds little to the critical path. The deployment also fixes defects in the original code, including end-token recovery and PAD-token handling, and vectorizes parts of decoding; see Appendix C for details.

## 6 Conclusion

We showed that temperature scaling provides a free lunch for autoregressive *de novo* peptide sequencing. Across Casanovo, InstaNovo, and ContraNovo, elevated decoding temperatures improve top-1 peptide recall without retraining, model updates, or additional decoding passes. Our implementation analysis separates two mechanisms: temperature can change candidate-pool composition, final candidate ranking, or both, depending on the scores used by each decoder. This distinction explains why the oracle increases with temperature for InstaNovo and ContraNovo but remains flat for Casanovo.

We then introduced FragR as a complementary, training-free method for recovering correct peptides that remain in the candidate pool but are not ranked first. It treats each pool as a spectrum-specific miniature retrieval space and combines the model score with b/y-ion ladder evidence and intensity-weighted fragment coverage computed from raw spectral peaks. The coefficients are frozen and shared across models, temperatures, and datasets. On two datasets derived from the nine-species benchmark—approximately 1.53 million monoisotopic spectra and 232,738 spectra including isotope-shifted precursors—combining a common decoding temperature of *T* = 1.8 with FragR improves peptide recall over the *T* = 1.0 baseline by 1.24–1.49 percentage points on V1. The largest macro-averaged combined improvement on the isotope-precursor subset is 6.69 percentage points. The asynchronous implementation agrees with offline rescoring candidate by candidate and adds little end-to-end inference latency. These results identify decoding temperature as a direct source of free accuracy gains and candidate selection as a complementary source of further improvement, without additional training or larger models.

## 7 Future Work

For all three models, the candidate-pool oracle exceeds the top-1 recall reported by RankNovo, although the rescored top-1 results do not yet reach that level. The remaining headroom motivates several directions for future work: developing PSM-scoring tools to rescore and rank candidate spectra directly; applying reranking models such as RankNovo to the top-10 beam candidates of a single model, avoiding the cost of multi-model inference; and exploring temperature scheduling strategies such as annealing, where the temperature varies across decoding steps instead of remaining fixed. More broadly, model- and task-aware temperature selection, information exchange among candidates, and generation-side pool expansion remain promising.

## Acknowledgments

This research was funded by Basic Research Program of Jiangsu. Grant number: BK20253035. This work was also supported by the Research Fund of Jiangsu Provincial Department of Science and Technology and Jiangsu Key Laboratory of AI for Industries.

## Code Availability

All three baseline models use their official releases. Casanovo uses the v4.0.0 source code ^1^ and the official weights trained on MassIVE-KB ^2^; v4.0.0 is the only release for which official MassIVE-KB-trained weights are available. InstaNovo uses the v1.2.0 source code ^3^ and the official v1.1.0 weights ^4^. ContraNovo uses the official source code ^5^ and official weights ^6^. The FragR implementation, including the asynchronous inference modifications to all three repositories, the rescoring code, and all analysis scripts, is available at https://github.com/sword-nan/FragR.

## Data Availability

The nine-species V1 dataset was obtained from MassIVE dataset MSV000081382 ^7^. The underlying V2 spectra come from nine-species-main.zip ^8^. As described in Section 5.1, we selected 77 complete MGF files at the file level to balance isotope-spectrum counts across species.

## Appendix A: Dataset and Species-Level Results

To balance isotope-spectrum counts across species, we selected 77 complete MGF files at the file level from nine-species-main.zip. Tables A1–A3 report the V1 spectrum count for each species, the V2 spectrum distribution, and the complete list of selected files, respectively. Tables A4 and A5 provide species-level recall results. Figure A1 reports recall@*k* curves before and after rescoring across decoding temperatures, and Figures A2 and A3 report the ablation of the evidence terms and the weight-grid sensitivity scan.

**Table A1:** Numbers of spectra by species in the nine-species V1 dataset.

| Species | Spectra |
|---|---|
| bacillus | 291,783 |
| honeybee | 314,571 |
| tomato | 290,050 |
| mmazei | 164,421 |
| clambacteria | 150,611 |
| human | 130,583 |
| yeast | 111,312 |
| ricebean | 37,775 |
| mouse | 37,021 |
| Total | 1,528,127 |

**Table A2:**
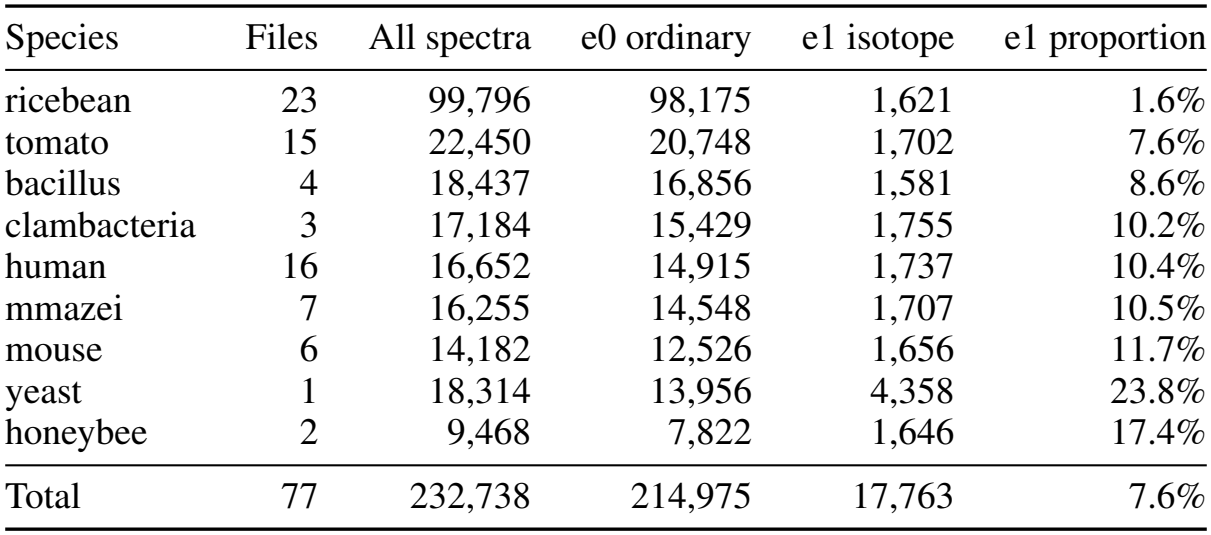
Spectrum distribution by species in the nine-species V2 dataset. Here, e0 denotes ordinary-precursor spectra and e1 denotes isotope-precursor spectra.

| Species | Files | All spectra | e0 ordinary | e1 isotope | e1 proportion |
|---|---|---|---|---|---|
| ricebean | 23 | 99,796 | 98,175 | 1,621 | 1.6% |
| tomato | 15 | 22,450 | 20,748 | 1,702 | 7.6% |
| bacillus | 4 | 18,437 | 16,856 | 1,581 | 8.6% |
| clambacteria | 3 | 17,184 | 15,429 | 1,755 | 10.2% |
| human | 16 | 16,652 | 14,915 | 1,737 | 10.4% |
| mmazei | 7 | 16,255 | 14,548 | 1,707 | 10.5% |
| mouse | 6 | 14,182 | 12,526 | 1,656 | 11.7% |
| yeast | 1 | 18,314 | 13,956 | 4,358 | 23.8% |
| honeybee | 2 | 9,468 | 7,822 | 1,646 | 17.4% |
| Total | 77 | 232,738 | 214,975 | 17,763 | 7.6% |

**Table A3:** Selected MGF files for the nine-species V2 dataset (77 files in total).

| Species | File |
|---|---|
| bacillus | 150723_QE1_PK_Bsub_Comp_PF_1.mgf |
|  | 150723_QE1_PK_Bsub_Exp_PF_1.mgf |
|  | 150723_QE1_PK_Bsub_Spore_PF_1.mgf |
|  | 151012_QEp_PK_Sec_6_DDA.mgf |
| clambacteria | Run1_82-g1_2000ng.mgf |
|  | Run2_82-g1_2000ng.mgf |
|  | Run2_83-g1_2000ng.mgf |
| honeybee | S-1.mgf |
|  | S-2.mgf |
| human | 151009_exo3_1.mgf |
|  | 151009_exo3_2.mgf |
|  | 151009_exo3_3.mgf |
|  | 151009_exo3_4.mgf |
|  | 151009_exo3_5.mgf |
|  | 151009_exo3_6.mgf |
|  | 151009_exo4_1_3h.mgf |
|  | 151009_exo4_4.mgf |
|  | 151009_exo4_4_3h.mgf |
|  | 151009_exo4_5.mgf |
|  | 151009_exo4_6.mgf |
|  | 151218_exo3_3.mgf |
|  | 151218_exo3_4.mgf |
|  | 151218_exo4_4.mgf |
|  | 151218_exo4_5.mgf |
|  | 151218_exo4_6.mgf |
| mmazei | Mm2DLC_N_2_01.mgf |
|  | Mm2DLC_N_2_09.mgf |
|  | Mm2DLC_N_2_12.mgf |
|  | MmGelfreeLC_N_1_01.mgf |
|  | MmGelfreeLC_N_1_05.mgf |
|  | MmGelfreeLC_N_1_10.mgf |
|  | MmGelfreeLC_N_1_11.mgf |
| mouse | 20160323_CoAN_CTRL2_3373.mgf |
|  | 20160323_CoAN_Deliti3_3380.mgf |
|  | 20160323_CoAN_WT1_3375.mgf |
|  | 20160323_CoAN_WT2_3376.mgf |
|  | 20160323_CoAN_WT3_3377.mgf |
|  | 20160329_CoAN_CTRL3_DR_3406.mgf |
| ricebean | 4705.mgf |
|  | 4706.mgf |
|  | 4707.mgf |
|  | 4709.mgf |
|  | 4710.mgf |
|  | 4711.mgf |
|  | 4713.mgf |
|  | 4714.mgf |
|  | 4715.mgf |
|  | 4717.mgf |
|  | 4718.mgf |
|  | 4719.mgf |
|  | 4721.mgf |
|  | 4722.mgf |
|  | 4723.mgf |
|  | 4726.mgf |
|  | 4728.mgf |
|  | 4730.mgf |
|  | 4731.mgf |
|  | 4732.mgf |
|  | 4734.mgf |
|  | 4740.mgf |
|  | 4741.mgf |
| tomato | 03022016_Clara_MP_Fraction_02.mgf |
|  | 03022016_Clara_MP_Fraction_03.mgf |
|  | 03022016_Clara_MP_Fraction_04.mgf |
|  | 04022016_Clara_SP_Fraction_02.mgf |
|  | 04022016_Clara_SP_Fraction_03.mgf |
|  | 04022016_Clara_SP_Fraction_04.mgf |
|  | 04022016_Clara_SP_Fraction_05.mgf |
|  | 04022016_Clara_SP_Fraction_06.mgf |
|  | 04022016_Clara_SP_Fraction_25.mgf |
|  | 04022016_Clara_SP_Fraction_26.mgf |
|  | 04022016_Clara_SP_Fraction_27.mgf |
|  | 04022016_Clara_SP_Fraction_28.mgf |
|  | 04022016_Clara_SP_Fraction_29.mgf |
|  | 04022016_Clara_SP_Fraction_30.mgf |
|  | 04022016_Clara_SP_Fraction_31.mgf |
| yeast | delPBP1_DB_heat_1.mgf |

**Table A4:**
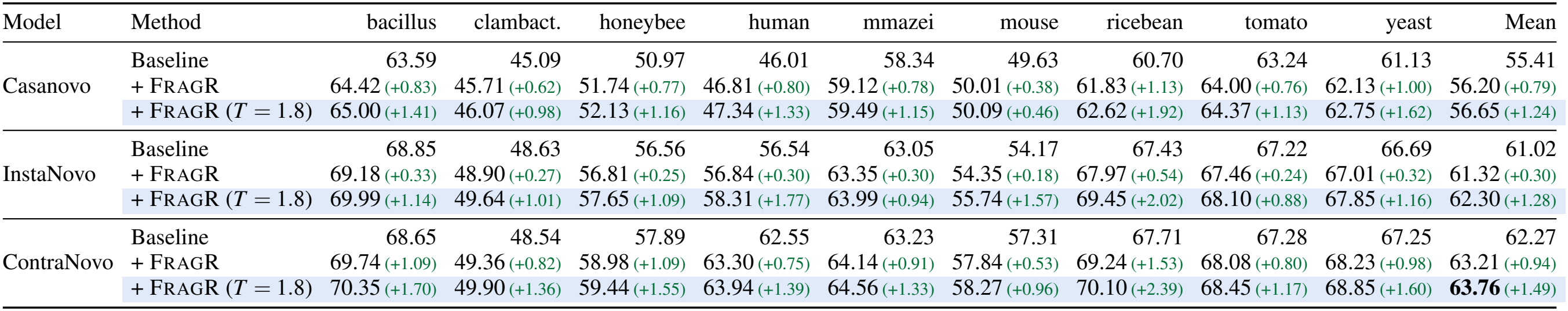
Species-level V1 peptide recall (%) and gains over the baseline for the same species, in percentage points. Light-blue rows combine rescoring with temperature scaling (*T* = 1.8). The best overall macro-average is bold; clambact. abbreviates clambacteria.

| Model | Method | bacillus | clambact. | honeybee | human | mmazei | mouse | ricebean | tomato | yeast | Mean |
|---|---|---|---|---|---|---|---|---|---|---|---|
| Casanovo | Baseline | 63.59 | 45.09 | 50.97 | 46.01 | 58.34 | 49.63 | 60.70 | 63.24 | 61.13 | 55.41 |
|  | + FragR | 64.42 (+0.83) | 45.71 (+0.62) | 51.74 (+0.77) | 46.81 (+0.80) | 59.12 (+0.78) | 50.01 (+0.38) | 61.83 (+1.13) | 64.00 (+0.76) | 62.13 (+1.00) | 56.20 (+0.79) |
|  | + FragR (*T* = 1.8) | 65.00 (+1.41) | 46.07 (+0.98) | 52.13 (+1.16) | 47.34 (+1.33) | 59.49 (+1.15) | 50.09 (+0.46) | 62.62 (+1.92) | 64.37 (+1.13) | 62.75 (+1.62) | 56.65 (+1.24) |
| InstaNovo | Baseline | 68.85 | 48.63 | 56.56 | 56.54 | 63.05 | 54.17 | 67.43 | 67.22 | 66.69 | 61.02 |
|  | + FragR | 69.18 (+0.33) | 48.90 (+0.27) | 56.81 (+0.25) | 56.84 (+0.30) | 63.35 (+0.30) | 54.35 (+0.18) | 67.97 (+0.54) | 67.46 (+0.24) | 67.01 (+0.32) | 61.32 (+0.30) |
|  | + FragR (*T* = 1.8) | 69.99 (+1.14) | 49.64 (+1.01) | 57.65 (+1.09) | 58.31 (+1.77) | 63.99 (+0.94) | 55.74 (+1.57) | 69.45 (+2.02) | 68.10 (+0.88) | 67.85 (+1.16) | 62.30 (+1.28) |
| ContraNovo | Baseline | 68.65 | 48.54 | 57.89 | 62.55 | 63.23 | 57.31 | 67.71 | 67.28 | 67.25 | 62.27 |
|  | + FragR | 69.74 (+1.09) | 49.36 (+0.82) | 58.98 (+1.09) | 63.30 (+0.75) | 64.14 (+0.91) | 57.84 (+0.53) | 69.24 (+1.53) | 68.08 (+0.80) | 68.23 (+0.98) | 63.21 (+0.94) |
|  | + FragR (*T* = 1.8) | 70.35 (+1.70) | 49.90 (+1.36) | 59.44 (+1.55) | 63.94 (+1.39) | 64.56 (+1.33) | 58.27 (+0.96) | 70.10 (+2.39) | 68.45 (+1.17) | 68.85 (+1.60) | **63.76** (+1.49) |

**Table A5:**
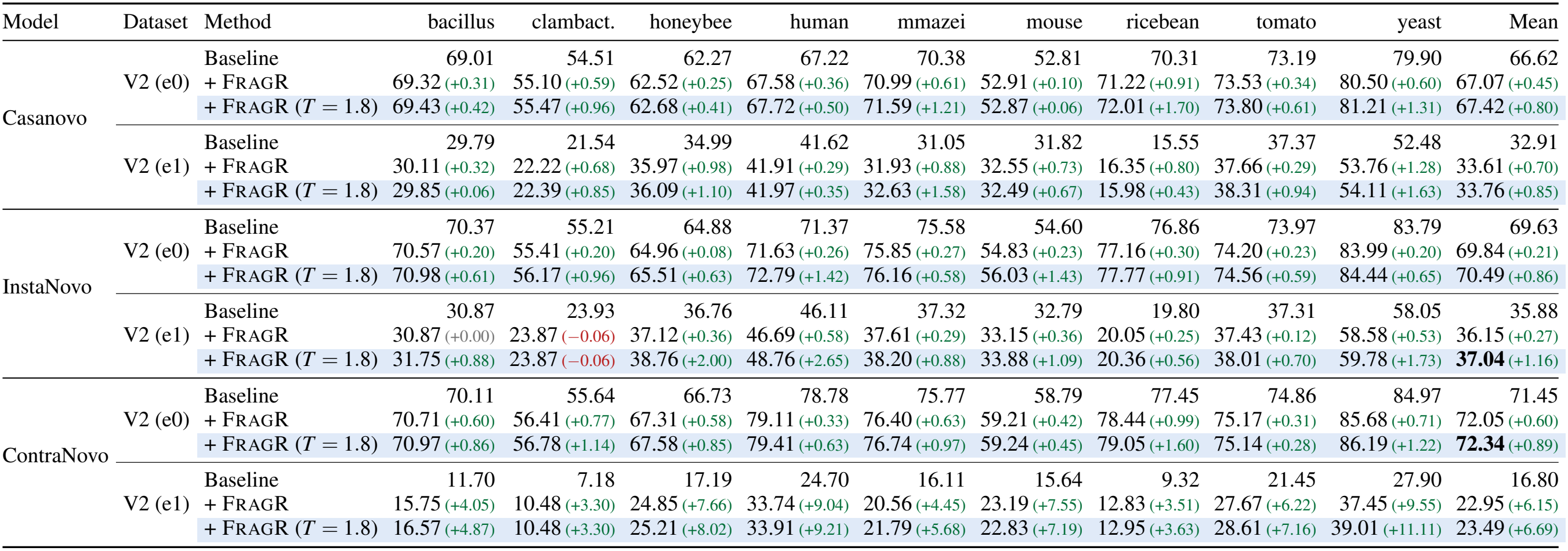
Species-level V2 peptide recall (%) and gains over the baseline for the same species, in percentage points. Here, e0 is the ordinary-precursor subset and e1 is the isotope-precursor subset. Light-blue rows combine rescoring with temperature scaling (*T* = 1.8), negative gains are shown in red, and the best macro-average in each subset is bold; clambact. abbreviates clambacteria.

| Model | Dataset | Method | bacillus | clambact. | honeybee | human | mmazei | mouse | ricebean | tomato | yeast | Mean |
|---|---|---|---|---|---|---|---|---|---|---|---|---|
| Casanovo | V2 (e0) | Baseline | 69.01 | 54.51 | 62.27 | 67.22 | 70.38 | 52.81 | 70.31 | 73.19 | 79.90 | 66.62 |
|  |  | + FragR | 69.32 (+0.31) | 55.10 (+0.59) | 62.52 (+0.25) | 67.58 (+0.36) | 70.99 (+0.61) | 52.91 (+0.10) | 71.22 (+0.91) | 73.53 (+0.34) | 80.50 (+0.60) | 67.07 (+0.45) |
|  |  | + FragR (*T* = 1.8) | 69.43 (+0.42) | 55.47 (+0.96) | 62.68 (+0.41) | 67.72 (+0.50) | 71.59 (+1.21) | 52.87 (+0.06) | 72.01 (+1.70) | 73.80 (+0.61) | 81.21 (+1.31) | 67.42 (+0.80) |
|  | V2 (e1) | Baseline | 29.79 | 21.54 | 34.99 | 41.62 | 31.05 | 31.82 | 15.55 | 37.37 | 52.48 | 32.91 |
|  |  | + FragR | 30.11 (+0.32) | 22.22 (+0.68) | 35.97 (+0.98) | 41.91 (+0.29) | 31.93 (+0.88) | 32.55 (+0.73) | 16.35 (+0.80) | 37.66 (+0.29) | 53.76 (+1.28) | 33.61 (+0.70) |
|  |  | + FragR (*T* = 1.8) | 29.85 (+0.06) | 22.39 (+0.85) | 36.09 (+1.10) | 41.97 (+0.35) | 32.63 (+1.58) | 32.49 (+0.67) | 15.98 (+0.43) | 38.31 (+0.94) | 54.11 (+1.63) | 33.76 (+0.85) |
| InstaNovo | V2 (e0) | Baseline | 70.37 | 55.21 | 64.88 | 71.37 | 75.58 | 54.60 | 76.86 | 73.97 | 83.79 | 69.63 |
|  |  | + FragR | 70.57 (+0.20) | 55.41 (+0.20) | 64.96 (+0.08) | 71.63 (+0.26) | 75.85 (+0.27) | 54.83 (+0.23) | 77.16 (+0.30) | 74.20 (+0.23) | 83.99 (+0.20) | 69.84 (+0.21) |
|  |  | + FragR (*T* = 1.8) | 70.98 (+0.61) | 56.17 (+0.96) | 65.51 (+0.63) | 72.79 (+1.42) | 76.16 (+0.58) | 56.03 (+1.43) | 77.77 (+0.91) | 74.56 (+0.59) | 84.44 (+0.65) | 70.49 (+0.86) |
|  | V2 (e1) | Baseline | 30.87 | 23.93 | 36.76 | 46.11 | 37.32 | 32.79 | 19.80 | 37.31 | 58.05 | 35.88 |
|  |  | + FragR | 30.87 (+0.00) | 23.87 (−0.06) | 37.12 (+0.36) | 46.69 (+0.58) | 37.61 (+0.29) | 33.15 (+0.36) | 20.05 (+0.25) | 37.43 (+0.12) | 58.58 (+0.53) | 36.15 (+0.27) |
|  |  | + FragR (*T* = 1.8) | 31.75 (+0.88) | 23.87 (−0.06) | 38.76 (+2.00) | 48.76 (+2.65) | 38.20 (+0.88) | 33.88 (+1.09) | 20.36 (+0.56) | 38.01 (+0.70) | 59.78 (+1.73) | **37.04** (+1.16) |
| ContraNovo | V2 (e0) | Baseline | 70.11 | 55.64 | 66.73 | 78.78 | 75.77 | 58.79 | 77.45 | 74.86 | 84.97 | 71.45 |
|  |  | + FragR | 70.71 (+0.60) | 56.41 (+0.77) | 67.31 (+0.58) | 79.11 (+0.33) | 76.40 (+0.63) | 59.21 (+0.42) | 78.44 (+0.99) | 75.17 (+0.31) | 85.68 (+0.71) | 72.05 (+0.60) |
|  |  | + FragR (*T* = 1.8) | 70.97 (+0.86) | 56.78 (+1.14) | 67.58 (+0.85) | 79.41 (+0.63) | 76.74 (+0.97) | 59.24 (+0.45) | 79.05 (+1.60) | 75.14 (+0.28) | 86.19 (+1.22) | **72.34** (+0.89) |
|  | V2 (e1) | Baseline | 11.70 | 7.18 | 17.19 | 24.70 | 16.11 | 15.64 | 9.32 | 21.45 | 27.90 | 16.80 |
|  |  | + FragR | 15.75 (+4.05) | 10.48 (+3.30) | 24.85 (+7.66) | 33.74 (+9.04) | 20.56 (+4.45) | 23.19 (+7.55) | 12.83 (+3.51) | 27.67 (+6.22) | 37.45 (+9.55) | 22.95 (+6.15) |
|  |  | + FragR (*T* = 1.8) | 16.57 (+4.87) | 10.48 (+3.30) | 25.21 (+8.02) | 33.91 (+9.21) | 21.79 (+5.68) | 22.83 (+7.19) | 12.95 (+3.63) | 28.61 (+7.16) | 39.01 (+11.11) | 23.49 (+6.69) |

## Appendix B: Decoding Scores and Temperature Scaling

### B.1 Notation and Sequence Scoring

Let *X* denote the encoded spectrum and *V* the vocabulary size. At decoding step *i*, the decoder output is projected through a prediction head to produce logits **l**^(*i*)^ ∈ ℝ^*V*^, which softmax converts to the probability vector **p**^(*i*)^ = Softmax(**l**^(*i*)^). The probability assigned to the selected token *w*_*i*_ is

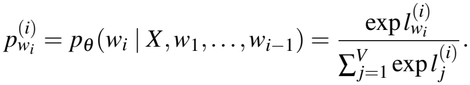

Its logarithm is 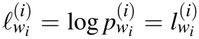 − logsumexp _*j*_ 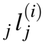. An autoregressive model defines the conditional sequence probability as 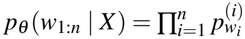, and beam search approximately maximizes its logarithm:

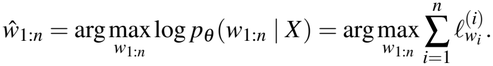

At each step, the search retains the *K* prefixes with the highest cumulative scores and expands them by one token. A final ranking score then selects the output sequence.

### B.2 Model-Specific Search and Ranking Scores

In practice, the score used during search need not be the same as the score used for final ranking, and the three implementations differ in this respect (Table A6).

**Table A6:** Search scores, final ranking scores, and the effect of temperature on the candidate pool for the three baseline models.

| Model | Search (top-*K* pruning) | Final ranking score | Temperature changes pool |
|---|---|---|---|
| Casanovo | Mean logits $\frac{1}{i}\sum_{t=1}^{i} l_{w_t}^{(t)}$ | Mean probabilities $\frac{1}{i}\sum_{t=1}^{i} p_{w_t}^{(t)}$ | No |
| InstaNovo | Cumulative log-probabilities $\sum_{t=1}^{i} \ell_{w_t}^{(t)}$ | Equivalent to the search score | Yes |
| ContraNovo | Mean probabilities $\frac{1}{i}\sum_{t=1}^{i} p_{w_t}^{(t)}$ | Original: mean after a second softmax; removed here (see B.3) | Yes |

Casanovo uses the stepwise mean of logits as its search score, omitting the LogSumExp normalization term at each step. Its final peptide score instead uses the mean of softmax probabilities. InstaNovo follows the standard procedure, accumulating log-probabilities after log_softmax without length normalization. ContraNovo uses mean probabilities during search. Its original implementation applies softmax again to each step’s probability vector during final ranking, giving 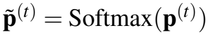 and the peptide score 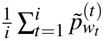.

### B.3 Aligning Model Scores for Rescoring

We made three modifications to place the three models’ scores on comparable scales. First, InstaNovo’s cumulative log-probability systematically favors shorter peptides because longer sequences naturally accumulate lower scores. We therefore use exp(score_raw*/L*) as its model score, where *L* is the number of candidate tokens. This is the geometric mean of token probabilities and, like the mean probabilities used by the other two models, lies in [0, 1]. Second, we remove ContraNovo’s second softmax during final ranking so that its ranking score matches its search score and reduces to the mean of ordinary softmax probabilities. Third, the constant −1 penalty that Casanovo and ContraNovo natively apply to precursor-mass-inconsistent candidates is removed from the model scores; mass consistency is instead handled by the penalty term in Eq. (2). All three modifications are applied to both the baseline and rescoring, so the comparison between them is unaffected. Throughout the paper, *baseline ranking* refers to these model-specific scores after alignment, not to InstaNovo’s unmodified native ranking.

### B.4 Model-Specific Effects of Temperature Scaling

For a temperature *T >* 0, the token distribution at decoding step *i* is

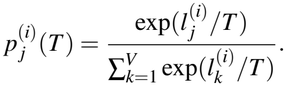

A larger *T* flattens the token distribution, whereas a smaller *T* sharpens it; *T* = 1 recovers the original softmax distribution [Hinton et al., 2015]. This statement concerns only the token probabilities. Whether temperature also changes the beam-search candidate pool depends on the scoring stage to which it is applied in each implementation.

After introducing temperature *T*, the affected stage differs across the three models. Casanovo applies temperature to logits. Because its search ranks candidates by mean logits, global scaling preserves their relative order and leaves the candidate-pool composition unchanged. Final scoring uses mean softmax probabilities, whose distribution changes with temperature; only the final scores therefore change. For InstaNovo, the accumulated log-probabilities change with temperature, affecting both the search process and the candidate pool. ContraNovo’s search directly uses mean probabilities, so temperature changes the probability distribution and likewise alters the candidate pool. This explains why Casanovo’s oracle curve in Fig. 3 is flat, whereas those of the other two models increase with temperature.

## Appendix C: Implementation and Inference Efficiency

### C.1 Repository-Specific Modifications and Optimizations

We made the necessary engineering modifications to the inference implementations in all three repositories. For defect correction, we fixed ContraNovo’s recovery of predictions at the end token and corrected InstaNovo’s handling of padding (PAD) tokens in the original code. For acceleration, Casanovo uses a vectorized beam-termination path, retains all completed candidates in the beam cache, and keeps top-*k* operations on the GPU throughout decoding. InstaNovo incorporates a fast decoder and vectorized knapsack decoding. ContraNovo uses a rewritten, vectorized beam-search implementation.

### C.2 Throughput and End-to-End Overhead

Figure A4 compares inference throughput before and after optimization, with all timings restricted to the inference loop (beam size 10, one RTX 4090; model loading excluded). In the synchronous pipeline, the loop covers model inference and exporting the prediction CSVs—top-1 results only, or the entire candidate pool when history dumping is enabled: each batch’s records are serialized and appended to disk before the next batch decodes. The asynchronous pipeline instead splits each batch after inference: the inference thread merely publishes the raw candidate tensors to shared memory, while a separate CPU process concurrently performs candidate decoding, fragment-feature computation, the precursor-mass veto, top-1 selection, and file writing.

On the nine-species V1 benchmark, optimization alone raises throughput for Casanovo, InstaNovo, and ContraNovo from 9.95, 6.39, and 3.32 to 56.57, 48.05, and 40.22 spectra/s, respectively—speedups of 5.68*×*, 7.52*×*, and 12.12*×*. Synchronously exporting the entire candidate pool inside the loop reduces throughput to 52.58, 46.58, and 38.61 spectra/s, a loss of 3–7%.

The reason synchronous export is costly, and asynchronous export is not, lies in when the CPU work is paid. In a synchronous loop the GPU stands idle whenever the inference thread performs CPU-side work—candidate bookkeeping, record serialization, file I/O—and enabling full-pool export lengthens exactly these segments, since collecting candidate tensors inside the loop forces frequent GPU synchronization. The asynchronous pipeline changes how time is accounted for, not how much work is done: the two stages run concurrently, so end-to-end time is governed by the slower stage rather than the sum. Per batch, decoding takes about 2.7 s while scoring and exporting takes about 0.2 s—a tenfold margin that keeps the CPU stage fully hidden behind the GPU.

The full-scale mouse runs (37,021 spectra, *T* = 1.0, batch 200) quantify this for all three models. For Casanovo, synchronous top-1 decoding spends 512.9 s on inference and 10.6 s on host-to-device transfer plus serializing and writing the top-1 CSV (523.5 s, 70.7 spectra/s); synchronous full-pool export raises inference itself to 538.1 s (+25.2 s,the in-loop candidate collection) and loop overhead to 16.7 s (+6.1 s of CSV I/O), totalling 554.8 s (66.7 spectra/s). Asynchronously, the same export plus complete rescoring—44 s of CPU work in the consumer process—overlaps with decoding: wall-clock is 521.8 s (70.9 spectra/s), on par with the synchronous top-1-only loop, and rescoring without export takes 512.3 s (72.3 spectra/s). The same ordering holds for the other two models: for ContraNovo, the asynchronous full pipeline finishes in 589.6 s versus 624.7 s synchronously (62.8 versus 59.3 spectra/s), on par with its synchronous top-1-only loop (588.6 s); for InstaNovo, asynchronous export is even 4.4% faster than the synchronous top-1-only loop (583.2 s versus 610.0 s; 63.5 versus 60.7 spectra/s). Run-to-run variation is about 1%.

Across all 27 full-scale runs, end-to-end wall-clock time exceeds the GPU inference time of the same run by only 0.5–4.5%. In short, the synchronous loop pays its CPU overhead on the GPU’s clock, whereas FragR pays it on a separate core: rescoring is nearly free and does not slow down inference, and exporting the entire candidate pool costs about one percentage point of wall-clock on top.

**Figure A1:**
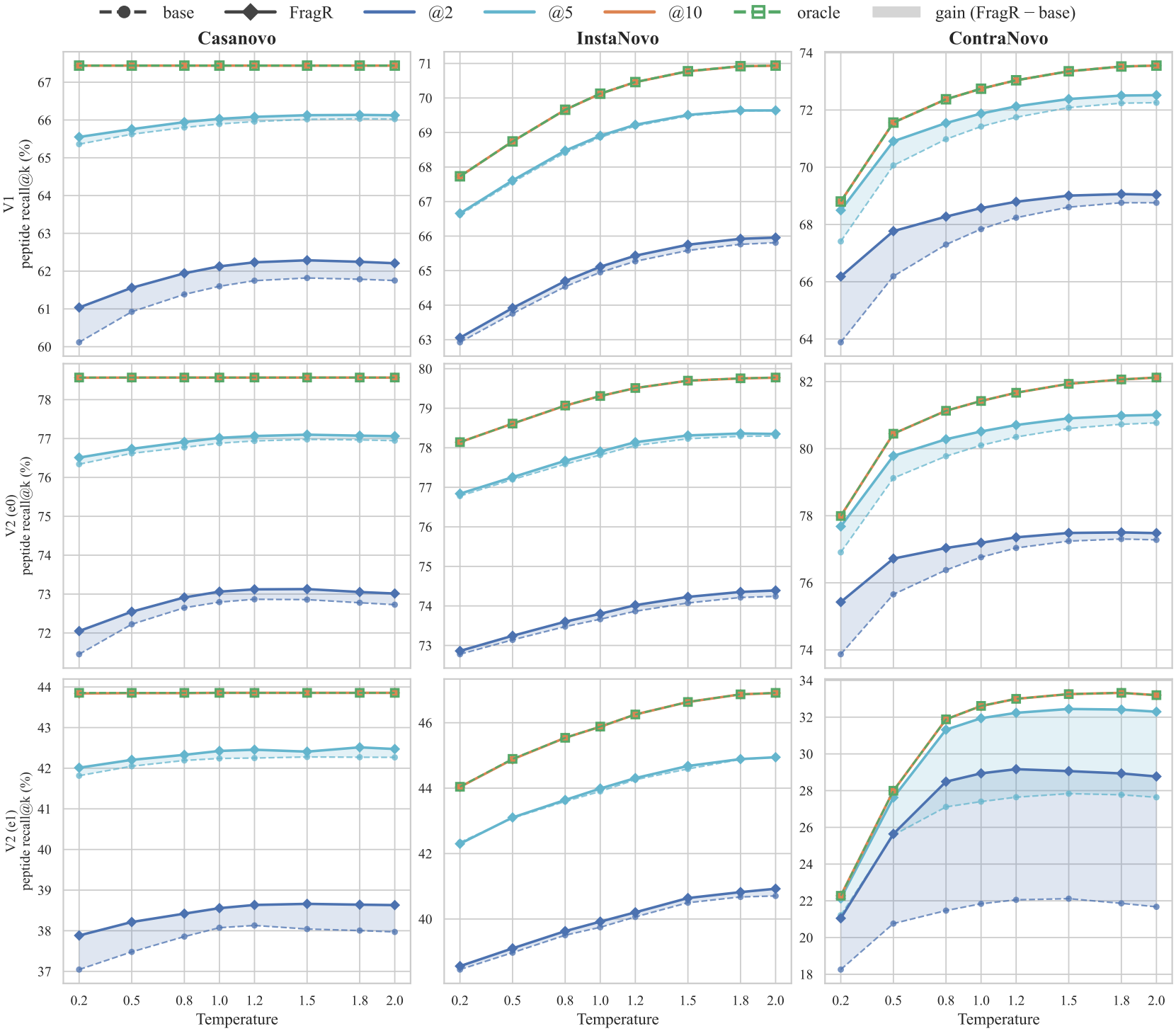
Recall@*k* (*k* = 2, 5, 10) before and after rescoring as a function of decoding temperature. In each upper panel, curves of the same color form a group: dashed lines denote baseline ranking and solid lines denote FragR ranking. The lower panels show their differences at the same cutoff depth, in percentage points. At *k* = 10, both rankings coincide with the oracle.

**Figure A2:**
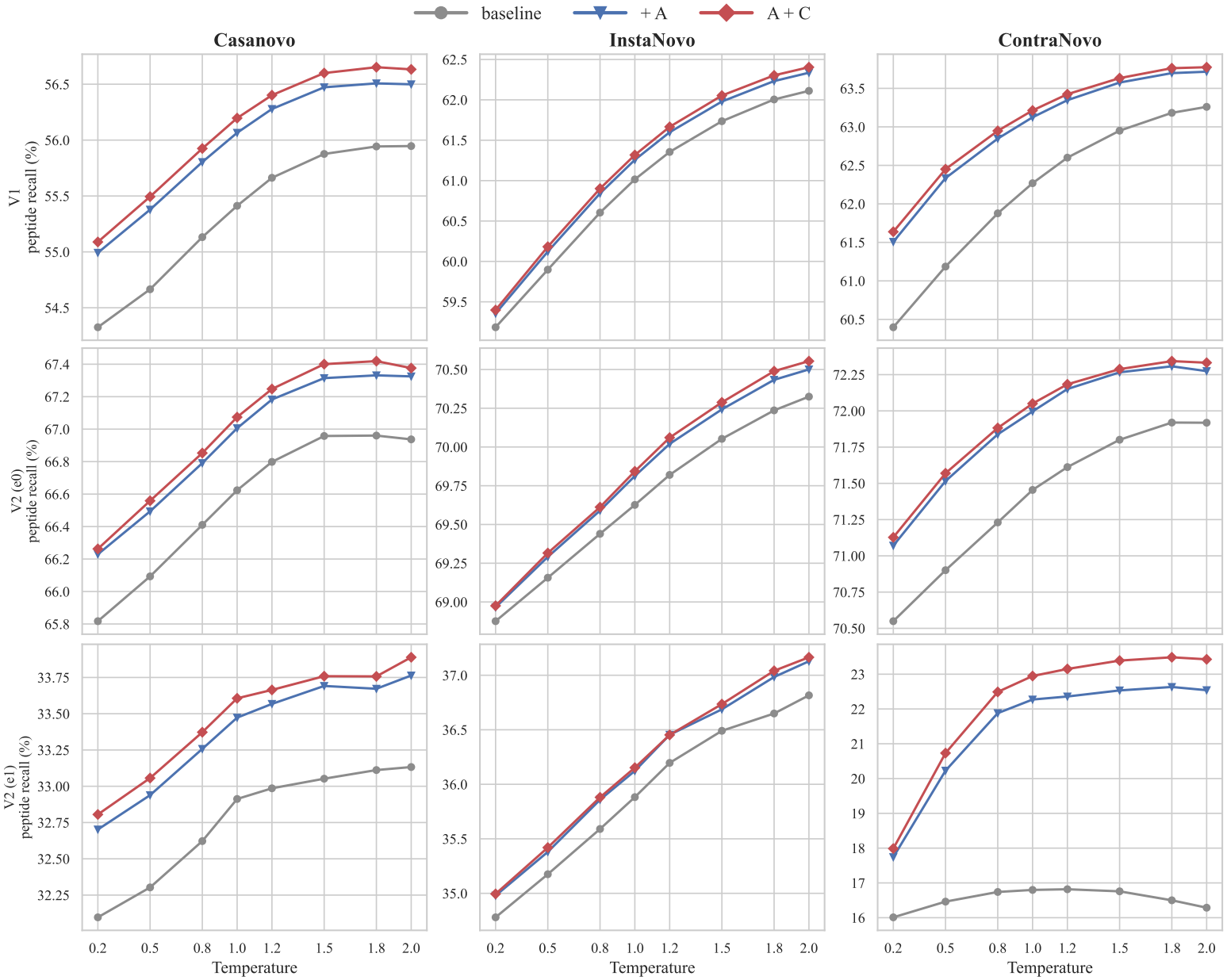
Ablation of the evidence terms: peptide recall as a function of temperature for the baseline, the ladder term *A* alone, and the complete score *A* + *C*. The *C*_*nl*_ label in the legend denotes *C* in the main text.

**Figure A3:**
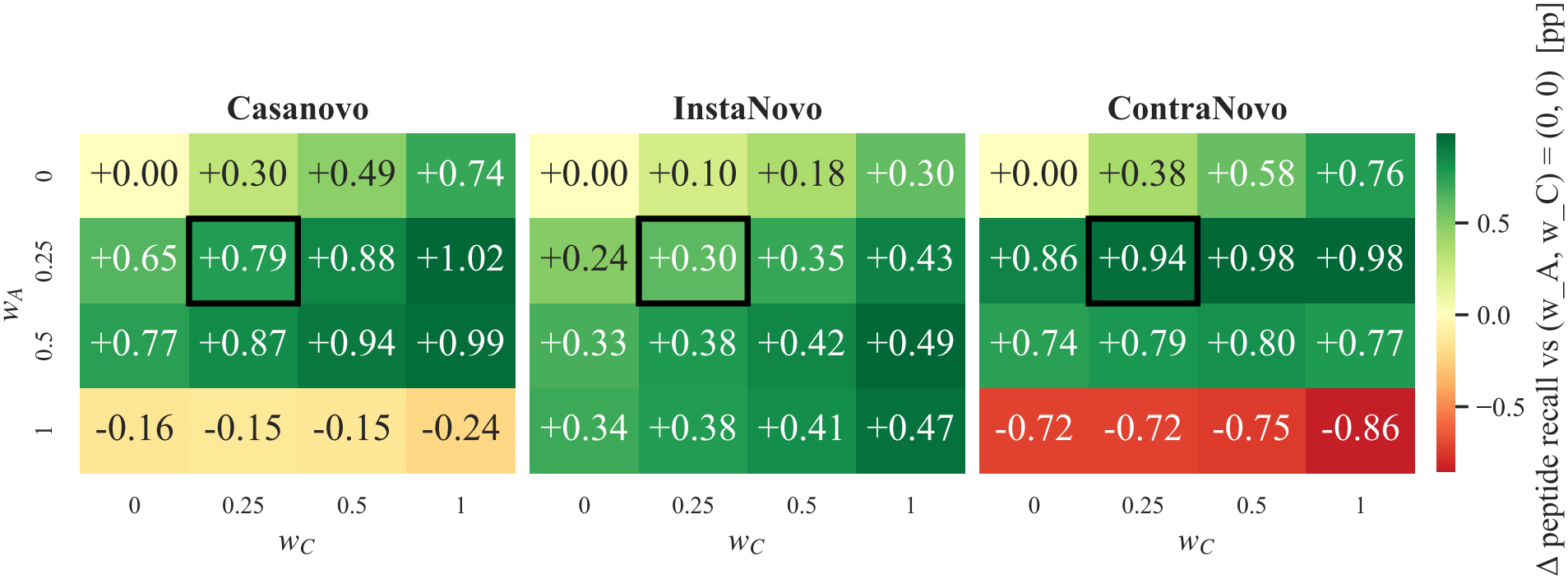
Weight-grid scan on the nine-species V1 dataset at *T* = 1.0: gains in peptide recall relative to (*w*_*A*_, *w*_*C*_) = (0, 0), in percentage points. The black box marks the frozen setting (0.25, 0.25).

**Figure A4:**
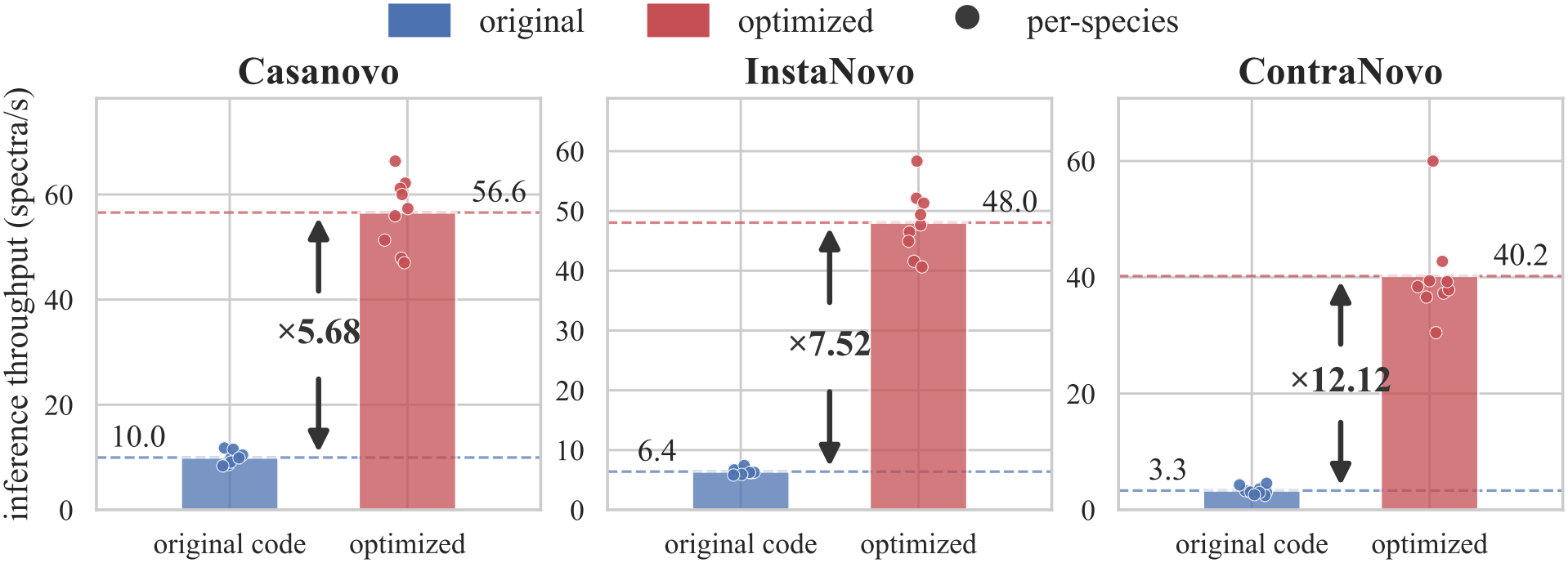
Inference throughput before and after engineering optimization on the nine-species V1 dataset, in spectra/s. Bars show macro-averages across the nine species, points show species-level measurements, and arrows annotate the speedup factors.

## Footnotes

1 https://github.com/Noble-Lab/casanovo/archive/refs/tags/v4.0.0.zip

2 https://github.com/Noble-Lab/casanovo/releases/download/v4.0.0/casanovo_massivekb.ckpt

3 https://github.com/instadeepai/InstaNovo/archive/refs/tags/1.2.0.zip

4 https://github.com/instadeepai/InstaNovo/releases/download/1.1.0/instanovo-v1.1.0.ckpt

5 https://github.com/BEAM-Labs/ContraNovo

6 https://drive.google.com/file/d/1knNUqSwPf98j388Ds2E6bG8tAXx8voWR/view

7 https://massive.ucsd.edu/ProteoSAFe/dataset.jsp?accession=MSV000081382

8 https://zenodo.org/records/12819175

